# Natural IgE promotes epithelial hyperplasia and inflammation-driven tumour growth

**DOI:** 10.1101/782805

**Authors:** Mark David Hayes, Sophie Ward, Greg Crawford, Rocio Castro Seoane, William David Jackson, David Kipling, David Voehringer, Deborah Dunn-Walters, Jessica Strid

**Author notes:** Corresponding author: Jessica Strid, Department of Immunology and Inflammation, Imperial College London, London, W12 0NN, UK.

## Abstract

IgE is the least abundant circulating antibody class but is constitutively present in healthy tissues bound to resident cells via its high-affinity receptor, FcεRI. The physiological role of endogenous IgE is unclear but it is suggested to provide host protection against a variety of noxious environmental substances and parasitic infections at epithelial barrier surfaces. Here we show that skin inflammation enhances levels of IgE with natural specificities and with a similar repertoire, VDJ rearrangements and CDRH3 characteristics as in healthy tissue. IgE-bearing basophils are recruited to inflamed skin via CXCL12 and TSLP/IL-3-dependent upregulation of CXCR4. In the inflamed skin, IgE/FcεRI-signalling in basophils promotes epithelial cell growth and differentiation, partly through histamine engagement of H_1_R and H_4_R. Furthermore, this natural IgE response strongly drives tumour outgrowth of epithelial cells harbouring oncogenic mutation. These findings indicate that natural IgE support skin barrier defences however during chronic tissue inflammation this may be subverted to promote tumour growth.

## Introduction

IgE is present at low concentrations in the blood and is the least abundant circulating antibody class. However, all epithelial tissues contain resident cells constitutively binding IgE antibodies. While the half-life of free IgE in the serum is short (1-2 days), tissue-resident IgE may persists for weeks or months_1_. In the tissue, IgE is bound with high affinity to its receptor, FcεRI, and can remain bound for the life of the FcεRI-bearing cell_2_. The specificity and function of this endogenous IgE in healthy individuals is unclear, although spontaneous production of natural IgE has been suggested to be a component of innate tissue defence_3, 4_.

Mast cells and basophils are the only two cell types that express the complete high-affinity receptor for IgE (FcεRI) with all four polypeptide subunits (αβγ2). In humans, subsets of dendritic cells, monocytes and eosinophils can express low levels of FcεRI but only as a trimer, lacking the β chain (αγ2). The density of mast cell and basophil FcεR1 expression correlates directly with serum IgE levels_5_. Mast cells and basophils are potent innate effector cells with overlapping effector functions. Paul Ehrlich first described mast cells and basophils in 1878 and estimated that there were enough mast cells in the body to make up ‘an organ the size of the spleen’6. Mast cells are constitutively resident in tissues, such as the skin, while basophils are circulating cells which rapidly infiltrate inflamed tissues. Crosslinking of FcεRI-bound IgE on these cells causes degranulation and immediate release of potent pharmacologically active pre-formed mediators and concurrent synthesis of cytokines/chemokines and inflammatory lipid mediators. Whilst the *in vivo* role(s) of both cell types is only partially understood, it has been suggested that they mediate a ‘gate-keeper’ function and promote barrier defences at skin and mucosal surfaces_7_.

IgE has most commonly been studied in the context of atopic allergic diseases or parasitic infections. However, it has been proposed to also play an important protective role in host defence against noxious environmental substances such as venoms, hematophagous fluids, environmental xenobiotics and irritants_4, 8_. In support of this, we recently demonstrated that topical exposure to environmental carcinogens promotes potent *de novo* induction of autoreactive IgE antibodies and that this IgE provides protection against epithelial carcinogenesis in the exposed tissue_9_. The induction of IgE was dependent on epithelial cell (EC) DNA-damage and deep-sequencing of the IgE antibodies revealed that they have a unique repertoire with specific VDJ rearrangements and CDRH3 characteristics distinct from IgE produced during general tissue inflammation_9_.

Epidermal hyperplasia and inflammation are hallmarks of a wide range of skin conditions. Furthermore, chronic inflammation may aid the proliferation and survival of EC harbouring genomic alterations and as such is now widely accepted to promote cancer development_10, 11, 12_. Here we explored the nature of inflammation-induced IgE antibodies and their role in tissue homeostasis. We show that general tissue inflammation enhances the levels of polyclonal natural IgE with similar characteristics and VDJ usage as in naïve animals. Furthermore, this natural IgE, via FcεRI-signalling in basophils, potently promotes epidermal hyperplasia and inflammation-driven outgrowth of skin tumours following subclinical carcinogen exposure. Tissue inflammation recruits IgE-bearing basophils in to the skin via CXCL12 and TSLP/IL-3-dependent upregulation of CXCR4 on circulating basophils. IgE-dependent basophil degranulation in the skin supports EC proliferation and differentiation, partly via histamine. Overall, our studies demonstrate how endogenous IgE with natural specificities potently affects skin EC growth and differentiation with implications for both atopy and cancer.

## Results

### Skin inflammation increases IgE levels locally and systemically

Both basophils and mast cells carry a high number of FcεRI receptors which are usually occupied with IgE_5_. Consistent with this, a horizontal view of resting healthy murine skin revealed large amounts of IgE normally present in the tissue (Fig. 1a) despite very low levels in serum (Fig. 1b-d). IgE in resting skin was mainly found on cells around the hair-follicles (Fig. 1a). Topical exposure to agents inducing skin inflammation, such as 12-*O*-Tetradecanoylphorbol-13-acetate (TPA, a protein kinase C activator) (Fig. 1b), MC903 (a vitamin D3 analogue commonly used to induce atopic dermatitis-like inflammation) (Fig. 1c) and R848 (Resiquimod, a toll-like receptor 7 agonist commonly used to induce psoriasis-like inflammation) (Fig. 1d) significantly enhanced the circulating levels of IgE compared to those in untreated (UT) or vehicle treated animals. Enhancement of serum IgE was dependent on topical exposure as IV or IP administration did not trigger the same effect, as shown by R848 exposure (Fig. 1d). The increase in serum IgE was accompanied by an increase in the number of IgE secreting plasma cells in the skin draining LNs (Fig 1e,f). IgE levels were also increased locally in the skin following topical TPA (Fig. 1g), but most notably the cells carrying the IgE switched from being predominantly mast cells in resting UT skin to mainly basophils in TPA-treated inflamed skin (Fig. 1h,i). 24h after cessation of TPA treatment only few mast cells remained in the skin, while IgE-bearing basophils accounted for ~2% of total CD45+ leukocytes. Basophil numbers further increased at 48h and then declined as the inflammation subsided, whilst mast cells returned to the skin (Fig. 1i). Thus, UT resting skin contains high levels of IgE which are predominately carried on mast cells. Skin inflammation increases IgE levels locally and systemically and IgE in inflamed skin is mainly carried on basophils.

**Figure 1.**
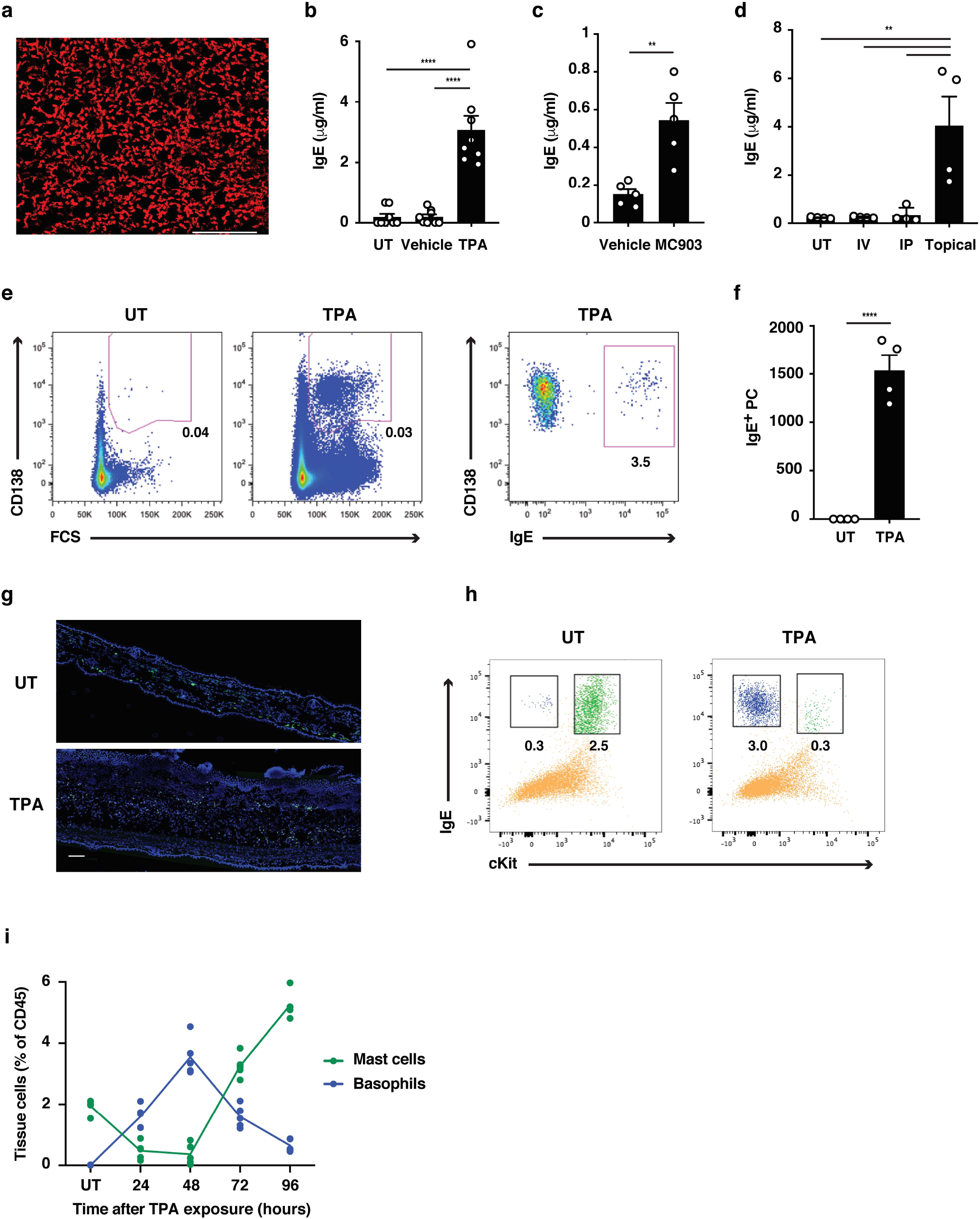
Skin inflammation increases IgE levels locally and systemically. (a) Representative image of IgE staining (red) in healthy dermal skin. Scale = 50μm. Image is representative of tile-scans from healthy UT dermal sheets of 5 independent mice. (b-d) ELISA of IgE in serum of UT WT BALB/c mice and mice treated on the dorsal ear skin with (b) 2.5nM TPA 2x a week for 2 weeks or vehicle control (ethanol) (n=8), (c) 1nM MC903 5x a week for 2 weeks or vehicle control (ethanol) (n=5) or (d) 100μg R848 3x a week for 2 weeks or similarly i.v. or i.p. (n=4). Data expressed as means ± SEM. (e,f) FACS analysis of IgE secreting plasma cells in the skin draining LNs of UT WT mice and mice exposed topically to TPA on the dorsal side of the ears 2x a week for 2 weeks. (e) Representative plots of plasma cells gated as FSC_hi_CD95_+_CD138_+_ cells, and with intracellular IgE staining to show isotype switching. (f) Enumeration of IgE_+_ plasma cells in skin draining LN (n=4). (g) Representative images of IgE staining (green) in UT and TPA treated whole skin. Nuclei in blue. Scale = 100μm. Images are representative of tile-scans from ear cross-sections from 6 independent mice. (h,i) FACS analysis of IgE-bearing cells in whole naïve UT skin and in TPA-treated skin (TPA 2x a week for 2 weeks). (h) Representative plots of IgE-bearing mast cells (green) and basophils (blue) 48h after last TPA exposure. (i) Enumeration of IgE-bearing mast cells and basophils at indicated time points after topical TPA. Mast cells were defined as CD45_hi_cKit_+_IgE_+_CD41_-_ and basophils as CD45_lo_cKit_-_IgE_+_ CD41_+_(n=5). Statistics using one-way ANOVA multiple comparison (b, d) and two-tailed Student’s t-test for unpaired data (c, f); **p<0.01 and ****p<0.0001. IV = intravenous; IP = Intraperitoneal.

### Skin inflammation enhances polyclonal ‘natural IgE’ with similar characteristics as in naïve animals

To understand the nature of the IgE induced during skin inflammation, we performed high-throughput sequencing of the IgE heavy-chain repertoire from mice exposed topically to TPA. This revealed the repertoires to be highly diverse and polyclonal with ~ 75 % of the repertoire consisting of clone sizes ≤ 10 (Fig. 2a), which was similar to that seen in UT naïve mice suggesting that no notable selection or clonal expansion had occurred. The most prominent effects of B cell selection can be seen in heavy-chain gene-usage and in CDRH3 characteristics. Thus we analysed the use of genes encoding the variable, diversity and joining (VDJ) regions of the IgE heavy-chain and found that IgHV1 was the most dominant Igh-V family gene accounting for > 70% of the repertoire (Fig. 2b). All 4 Igh-J family genes were used and 6 Igh-D family genes, although IgHD2 dominated (Fig. 2c). The overall VDJ family gene usage in UT naïve mice and in the TPA-treated mice was similar (Fig. 2b,c). Further, we analysed characteristics and physicochemical properties of the complementarity-determining CDRH3 regions as these form an important part of the antigen-binding site of the antibody. We found no differences in CDRH3 length, levels of mutation rate, aliphatic index or isoelectric point (pI) between the IgE sequenced from TPA-treated or UT mice (Fig. 2d), suggesting that the TPA-induced skin inflammation did not alter the composition or the nature of the IgE response. The dominant use of the V1 family and the low mutation rate is consistent with that of ‘natural IgE’3, thus we next tested whether the inflammation-induced IgE repertoire contained the characteristic natural anti-phosphorylcholine (anti-PC)_13_ specificity. This analysis showed that skin inflammation enhanced production of anti-PC IgE to a level similar to the spontaneous production of natural IgE seen in immunodeficient *Tcrb*_*−/−*_ mice (Fig. 2e). Levels of circulating anti-PC IgE increased further in skin tumour bearing mice that had been exposed to TPA for 20 weeks (Fig. 2e). Together this analysis revealed that skin inflammation enhances levels of undiversified natural IgE akin to the IgE spontaneously present at low levels in healthy mice.

**Figure 2.**
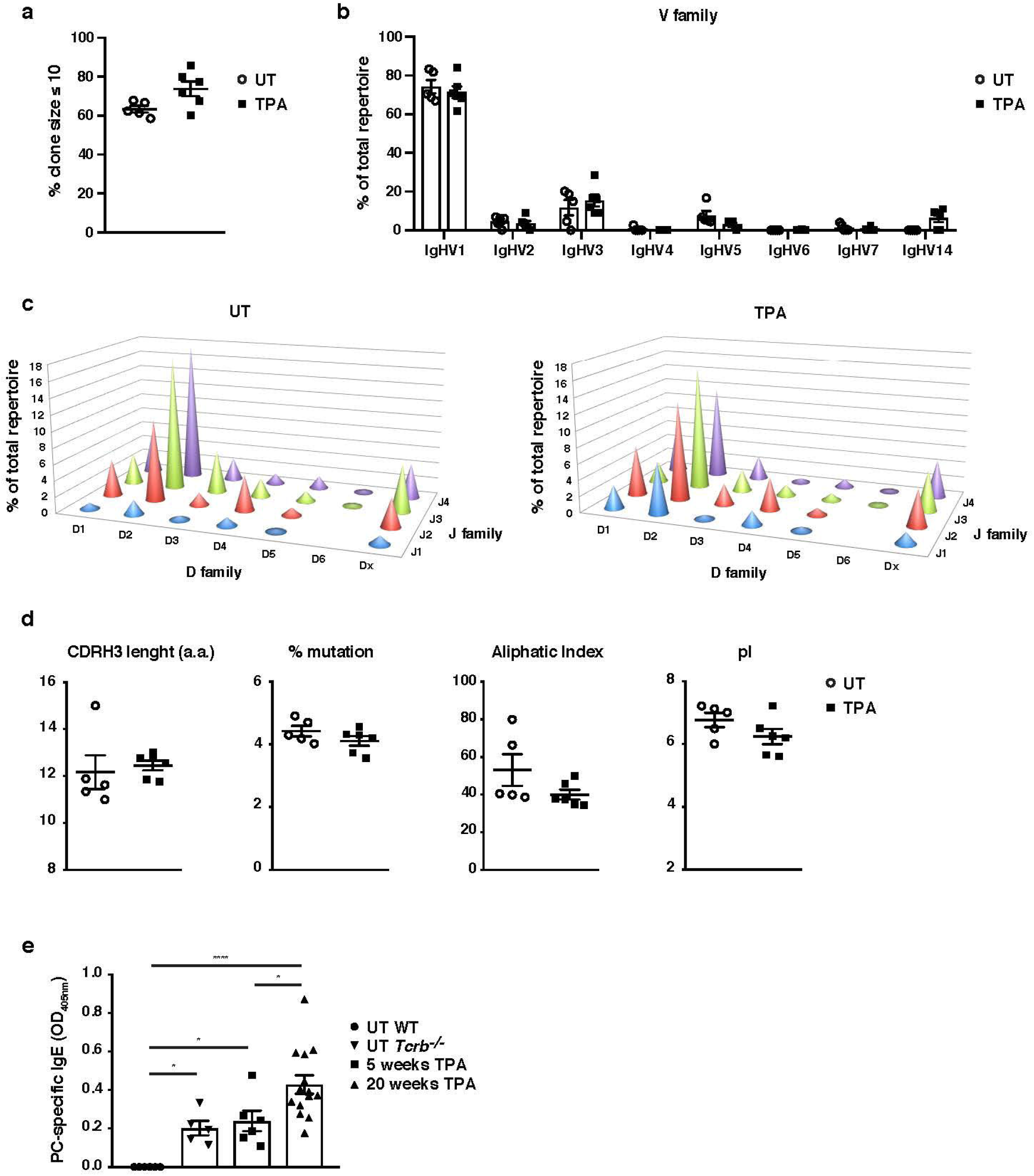
Skin inflammation induces polyclonal ‘natural IgE’ with similar characteristics to that in naïve animals. (a-d) High-throughput sequencing and heavy-chain repertoire analysis of IgE in sorted FSC_hi_CD95_hi_CD138_+_ plasma cells from skin draining LNs of WT mice 7 days after topical exposures to TPA (TPA 2x a week for 2 weeks) (n=6) and in whole spleen from UT naïve mice (n=5). (a) % of the total repertoire consisting of clone sizes ≤ 10. (b) Igh-V gene family usage expressed as mean of total repertoire ± SEM. V genes not used in the repertoire of any mouse removed for clarity. (b) Paired DJ repertoire shown as average Igh-D and Igh-J use of total repertoire in UT (left) and TPA-treated (right) mice. (d) Physicochemical properties of the CDRH3 regions; length (number of a.a.), % mutation compared to germline, aliphatic index and isoelectric point (pI) presented as mean ± SEM. (e) Level of IgE in serum specific for phosphorylcholine (PC) in UT WT (n=6), UT *Tcrb*_*−/−*_ (n=5), 10x TPA-treated (2x/week) WT mice (n=6) and in WT mice undergone DMBA-TPA carcinogenesis (single low-dose DMBA + 20 weeks TPA, 2x/week) (n=14). Data presented as means ± SEM. Statistics by two-tailed Student’s t-test for unpaired data (a, b and d) and one-way ANOVA multiple comparison (e); *p<0.05, and ****p<0.0001.

### Natural IgE promotes inflammation-driven outgrowth of tumours

We have recently reported, that IgE induced *de novo* following repeated topical exposure to environmental carcinogens is protective in a mutation-driven cutaneous carcinogenesis model and that the repertoire of the carcinogen-induced IgE differed substantially from the IgE induced by general skin inflammation_9_. We therefore next tested whether natural IgE would influence cutaneous carcinogenesis in an inflammation (TPA)-driven model. This was explored in a widely used two-stage model, in which topical exposure to a subclinal dose of carcinogen provokes few oncogenic mutations in EC (not enough for cancer growth) that can subsequently be promoted to grow into overt tumours by chronic tissue inflammation_14_. Mice lacking IgE (*Igh7*_*−/−*_) were markedly protected from developing tumours in this two-stage model (Fig. 3a). Whilst *Igh7*_*−/−*_ mice showed an equivalent level of DNA-damage, as assessed by staining for the phosphorylated histone H2A variant H2AX (γH2AX), to WT following carcinogen exposure (**Supp Fig. 1a**), only few *Igh7*_*−/−*_ mice developed tumours and the ones that did only grew very few and significantly smaller papillomas (Fig. 3a), suggesting that the lack of IgE during TPA promotion hindered tumour growth. The topical TPA application enhanced the circulating levels of IgE in WT animals throughout the experiment (Fig. 3b). *Igh7*_*−/−*_ mice developed no IgE and reduced levels of IgG1 and IgG2a compared to WT animals (**Supp Fig. 1b**). Analysis of the tumour tissue and the peri-lesional skin by flow cytometry (Fig. 3c) and by qRT-PCR (Fig. 3d) showed that basophils were the predominant cells carrying IgE in the tumours with very few mast cells entering the tumour. The peri-lesional skin contained both mast cells and basophils, whilst mast cells dominated in UT belly skin from the same animals (Fig. 3c,d). Cross-sections of whole tumours showed that IgE-bearing cells accumulated right up into the tumour, mainly in the peritumoural infiltrate (Fig. 3e), with some also entering the epithelium (Fig. 3e **inset**). Thus, TPA-enhanced natural IgE potently promotes the outgrowth of inflammation-driven tumours with IgE-bearing basophils accumulating inside skin tumours.

**Figure 3.**
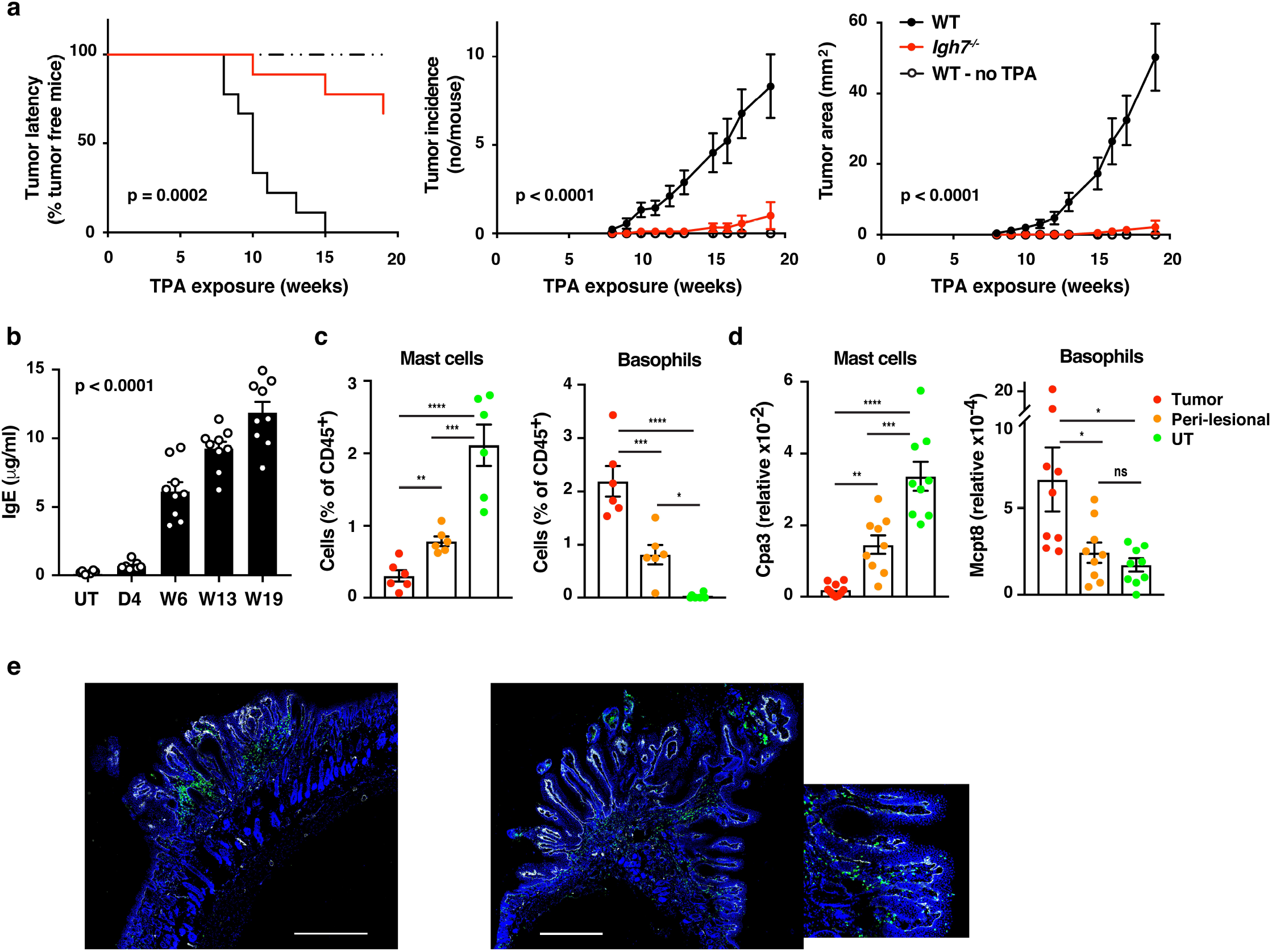
Natural IgE promotes inflammation-driven outgrowth of tumours. (a) Tumor susceptibility expressed as tumor latency (time to appearance of first tumor), tumor incidence (average number of tumors per mouse) and tumor area (average tumor size per mouse) in BALB/c WT and *Igh7*_*−/−*_ mice (n=9/group) mice following DMBA-TPA inflammation-driven carcinogenesis and following similar DMBA exposure without TPA (n=6). Data are expressed as means ± SEM and statistical significance assessed using Log-rank (Mantel-Cox) test for tumor latency and linear regression for tumor incidence and area. (b) ELISA of IgE in serum of WT mice at indicated time-points during DMBA-TPA carcinogenesis (n=9). (c) FACS analysis of IgE-bearing cells among the total CD45+ leukocyte infiltrate in tumour tissue, peri-lesional skin and UT belly skin of WT mice (n=6, tumours > 3mm were picked). Mast cells were defined as CD45_hi_cKit_+_IgE_+_CD41_-_ and basophils as CD45_lo_cKit_-_IgE_+_CD41_+_. (d) Quantitative RT-PCR analysis of Cpa3 transcripts (relative specific for mast cells) and Mcpt8 (relative specific for basophils) in tumour tissue, peri-lesional skin and UT belly skin from WT mice (n=9). Data are expressed as mean ± SEM relative to the control gene cyclophilin. (e) Representative images of IgE staining (green) and basement membrane component CD49F (white) in inflammation (DMBA-TPA)-induced tumours. Nuclei in blue. Scale = 400μm. Images are representative tile-scans of cross-sections from independent tumors in 8 WT mice. Inset shows higher magnification of indicated area. Statistics by one-way ANOVA testing for linear trend of IgE increase with time (b) and one-way ANOVA multiple comparison (c,d); *p<0.05, **p<0.01, ***p<0.001 and ****p<0.0001. ns = not significant.

### IgE exacerbates skin inflammation by altering the microenvironment and inducing epithelial hyperplasia

To further explore the role of IgE during TPA-induced inflammation (the tumour promotion phase), we investigated the skin microenvironment following repeated topical TPA application to the ear skin. This showed that TPA caused substantial *de novo* infiltration of CD45_+_ leukocytes in to the skin, which was equal in WT and *Igh7*_*−/−*_ mice (Fig. 4a). There was no substantial difference in the cellular composition of the infiltrate and importantly no difference in mast cell or basophil numbers either in UT or TPA-treated skin (Fig. 3a and Supp Fig. 2a). Similar results were found in TPA-treated *FceR1a*_*−/−*_ mice (**Supp Fig 2b**). Nevertheless, qRT-PCR analysis of whole skin during TPA-induced inflammation showed that IgE-deficient mice had significantly lower levels of IL-4, IL-5, IL-6 and IL-33 whilst producing similar levels of TNFα and higher levels of IL-25 (Fig. 4b). Furthermore, IgE-deficient animals showed no enhancement of the enzyme histidine decarboxylase (HDC) responsible for generating histamine and significantly lower levels of the enzymes involved in the production of bioactive prostanoids such as PGD_2_ and PGE_2_ compared to WT mice (Fig. 4b). Consistent with this, a significant amount of histamine was found to be released in the inflamed skin (Fig. 4c) and in to the serum (Fig. 4d) of TPA-treated WT mice, but not in in TPA-treated *Igh7*_*−/−*_ mice (Fig. 4c,d), suggesting that degranulation and histamine discharge was dependent on IgE engagement. Skin analysis by immunohistochemistry also showed obvious changes in the TPA-treated inflamed skin in the absence of IgE. Whilst both WT and *Igh7*_*−/−*_ mice showed a large and equal immune infiltrate in the dermis (Fig. 1a,e), significantly less epithelial hyperplasia was detected in the absence of IgE (Fig. 4e,f). Together these data demonstrate that the lack of IgE substantially alters the skin microenvironment and decreases inflammation-associated epithelial hyperplasia.

**Figure 4.**
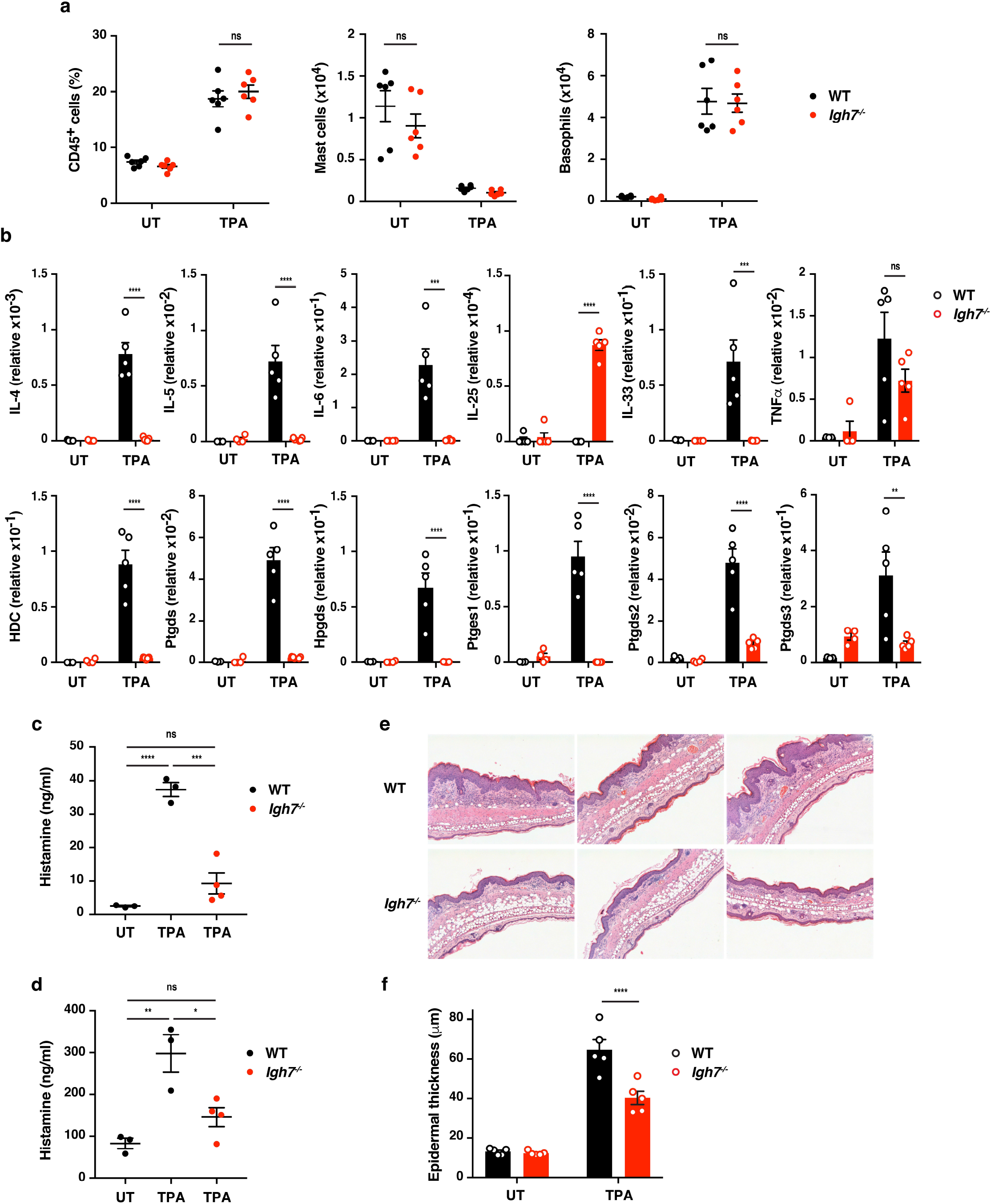
IgE alters the inflamed skin microenvironment and induces epithelial hyperplasia. (a-f) Analysis of the skin microenvironment in UT or TPA-treated (topical TPA to the dorsal ear skin 2x/week for 2 weeks) WT and *Igh7*_*−/−*_ mice. (a) FACS analysis of total CD45_+_ leukocytes in whole skin and enumeration of mast cells and basophil numbers (n=6). Mast cells were defined as CD45_hi_cKit_+_FcεRI_+_CD41_-_ and basophils as CD45_lo_cKit_-_FcεRI_+_CD41_+_. (b) Quantitative RT-PCR analysis of indicated transcripts in whole skin relative to the control gene cyclophilin (n=5). (c,d) ELISA assessing amount of histamine released *ex vivo* from ear skin in to media when floated overnight (c) and *in vivo* in to the serum (d) (WT n=3, *Igh7*_*−/−*_ n=4). (e) Representative images of H&E stained cross-sections from inflamed TPA-treated ear skin of WT and *Igh7*_*−/−*_ mice. (f) Epidermal thickness enumerated. Data in (a,b,c,d and f) are expressed as means ± SEM. Statistics by one-way ANOVA multiple comparison; *p<0.05, **p<0.01, ***p<0.001 and ****p<0.0001. ns = not significant.

### IgE activity in inflamed skin is mediated by FcεRI-signalling in basophils

To determine which cells were primarily responsible for the IgE-mediated exacerbation of skin inflammation, we sorted IgE-bearing mast cells and basophils from TPA-treated skin at a time-point when both cell populations were present. This showed that basophils produced vast amounts of cytokine such as IL-4, IL-6 and IL-13 and they were significantly more immune active than mast cells from the same tissue (Fig. 5a). Basophils sorted from the spleen of the same animal did not produce cytokines (Fig. 5b), suggesting that the basophils only became activated when entering the inflamed skin. IgE was partly required to activate the basophils for cytokine production when entering the skin (Fig. 5a) and IgE-signalling was necessary for degranulation and histamine release as histamine was absent from the skin and serum of *FceR1a*_*−/−*_ mice (Fig. 5c) as in *Igh7*_*−/−*_ mice (Fig. 4c,d). Indeed, mice with a constitutive depletion of basophils due to Cre toxicity_15_ (*Mcpt8*_*Cre/+*_) showed no release of histamine in inflamed skin or serum (Fig. 5c) and also demonstrated significantly less TPA-induced epithelial hyperplasia compared to WT mice (Fig. 5d). To investigate how IgE-mediated activation of basophils may affect skin EC, neonatal skin EC (keratinocytes) were grown *in vitro* and at 70% confluency supplemented with media from IgE-activated basophils or media alone as control. Basophil-derived conditioned media induced significant upregulation of Ki67 in EC, downregulation of the basal keratins K5 and K14 and upregulation of the suprabasal keratins K1 and K10 (Fig. 5e), suggesting that basophil-released mediators promoted proliferation and upwards differentiation of skin EC. Basophil media also induced expression of inflammatory cytokines such as IL-1α, IL-18 and IL-31 in EC (**Supp Fig. 3**). A similar pattern of induced expression of Ki67 and suprabasal keratin K1 and reduced expression of K14 was demonstrated when EC were grown *in vitro* with histamine (Fig. 5f) and the dose-dependent response to histamine was completely blocked when the EC cultures were pre-incubated with histamine receptor 1 or 4 (H_1_R or H_4_R) antagonists (Fig. 5f). Indeed, the basophil-induced EC proliferation and differentiation was also partially impeded by H_1_R or H_4_R blockade both *in vitro* (Fig. 5g) and *in vivo* (Fig. 5h). Thus, IgE-signalling in skin infiltrating basophils directly promote EC activation, proliferation and differentiation, partly via histamine receptors H_1_R and/or H_4_R on EC.

**Figure 5.**
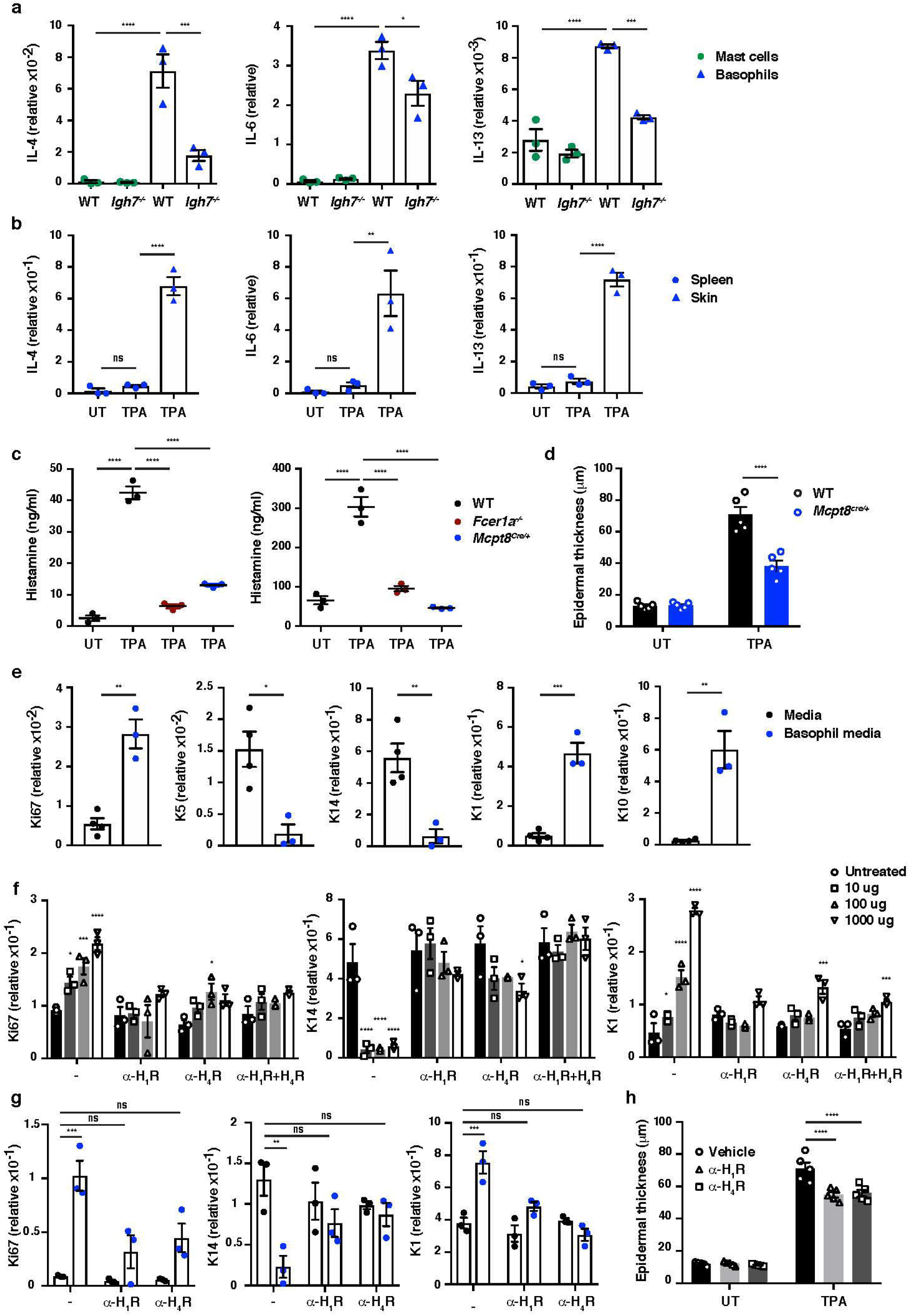
IgE activity in inflamed skin is mediated via FcεRI-signalling in basophils. (a-d,h) Healthy UT ear skin of mice was examined and compared to inflamed TPA-treated skin (topical TPA 2x/week for 2 weeks). (a) Mast cells and basophils were FACS sorted from the skin of WT and *Igh7*_*−/−*_ mice and (b) basophils were sorted from the spleen of WT mice 72h after the last TPA treatment (n=3/group). Mast cells were defined as CD45_hi_cKit_+_FcεRI_+_CD41_-_ and basophils as CD45_lo_cKit_-_FcεRI_+_CD41_+_. (a,b) Quantitative RT-PCR analysis of *Il4*, *Il6* and *Il13* transcripts in the FACS sorted cell populations from indicated tissue relative to the control gene cyclophilin. (c) ELISA assessing amount of histamine released *ex vivo* from ear skin in to media when floated overnight (left panel) and *in vivo* in to the serum (right panel) of WT, *FceR1a*_*−/−*_ and *Mcpt8*_*Cre/+*_ mice (n=3/group). (d) Epidermal thickness measured from H&E stained cross-sections of WT and *Mcpt8*_*Cre/+*_ ear skin (n=5/group). (e-g) Neonatal WT skin EC (keratinocytes) were grown *in vitro* to 70% confluency and then supplemented with (e) media from IgE-crosslinked basophils or media alone or (f) differing concentrations of histamine (n=3/group). Blocking of histamine receptors on ECs by adding 10μM H_1_R-or H_4_R-antagonist or both during cultures with added histamine (f) or basophil media (g) was also performed (n=3/group). (e-g) EC were analysed by qRT-PCR for expression of transcripts indicating cell cycling (Ki67) and differentiation (K5, K14, K1 and K10) relative to the control gene cyclophilin. (h) Epidermal thickness measured from H&E stained cross-sections of WT ear skin following *in vivo* blocking of H_1_R or H_4_R by i.p administration of antagonist drug or vehicle control (n=5/group). All data are expressed as means ± SEM. Statistics by one-way ANOVA multiple comparison (a-d, f-h) or two-tailed Student’s t-test (e); *p<0.05, **p<0.01, ***p<0.001 and ****p<0.0001. ns = not significant.

### Basophil recruitment to the skin is dependent on TSLP/IL-3-mediated upregulation of CXCR4

Next, we explored how IgE-bearing basophils are recruited to inflamed skin. Treating mice with pertussis toxin (PTX) prior to topical TPA significantly reduced the recruitment of basophils to the skin (Fig. 6a), indicating a role for G-protein coupled receptors such as chemokine receptors (CCRs). In support of this idea, TPA-treated inflamed skin showed increased levels of mRNA for *CCL2*, *CCL5*, *CCL8*, *CCL11* and *CXCL12* compared with normal UT skin (Fig. 6b). In parallel, infiltrating basophils showed preferential expression of *CXCR2*, *CXCR4* and *CrTH2* (PGD_2_ receptor) which differed significantly to the CCRs expressed on mast cells from the same tissue (Fig. 6c). Therefore, we tested the effect of blocking the most abundant chemokine receptors on basophil recruitment to inflamed skin. Blocking CXCR2 with a neutralising antibody inhibited the TPA-induced recruitment of neutrophils but not of basophils (Fig. 6d), while mice lacking CCR2 (*CCR2*_*−/−*_) had reduced recruitment of monocytes but not of basophils (Fig. 6e). Inhibition of CrTH2-signalling by COX-2 blockade or by injection of a small molecule inhibitor (AMG 853) had no effect on the recruitment of any cell type to the inflamed skin (data not shown). In contrast, blocking CXCR4 with a neutralising antibody significantly and selectively reduced basophil infiltration to the inflamed skin (Fig. 6f), suggesting a role for CXCL12-CXCR4 in driving the basophil recruitment. However, although resting basophils abundantly expressed CXCR4, most of this was intracellular (Fig. 6g); therefore we explored whether any mediators released during skin inflammation might upregulate surface expression of CXCR4 by basophils. Indeed, basophils stimulated with TSLP or IL-3, which are abundantly expressed in inflamed skin_16_, or PMA/ionomycin as a control, upregulated expression of CXCR4 on the cell surface (Fig. 6h-j). Both *de novo* expression of CXCR4 RNA was induced by TSLP (Fig. 6h) and the intracellular pool of CXCR4 protein was transported to the cell surface when basophils were stimulated with TSLP or IL-3 (**Fig. i,j**). As IL-3 is classically produced by T cells, we next examined the effect of depleting T cells and found a complete lack of basophil recruitment to the inflamed skin of mice lacking all T cells (Fig. 6k). In support, we found that reducing IL-3 levels during skin inflammation by intradermal injection of neutralising antibodies also reduced the recruitment of basophils (Fig. 6l). Similarly, blocking TSLP, significantly reduced basophil recruitment to inflamed skin and blocking TSLP and IL-3 simultaneously synergistically abolished basophil recruitment (Fig. 6l). Injection of blocking antibodies intradermally in one ear also blocked recruitment into the inflamed skin in the contralateral ear (Fig. 6m), suggesting that the cytokine(s) were acting systemically and not locally on basophil recruitment. Together this shows that production of TSLP and IL-3 in inflamed skin drives surface expression of CXCR4 on systemic basophils, allowing recruitment to the skin in response to increased levels of CXCL12.

**Figure 6.**
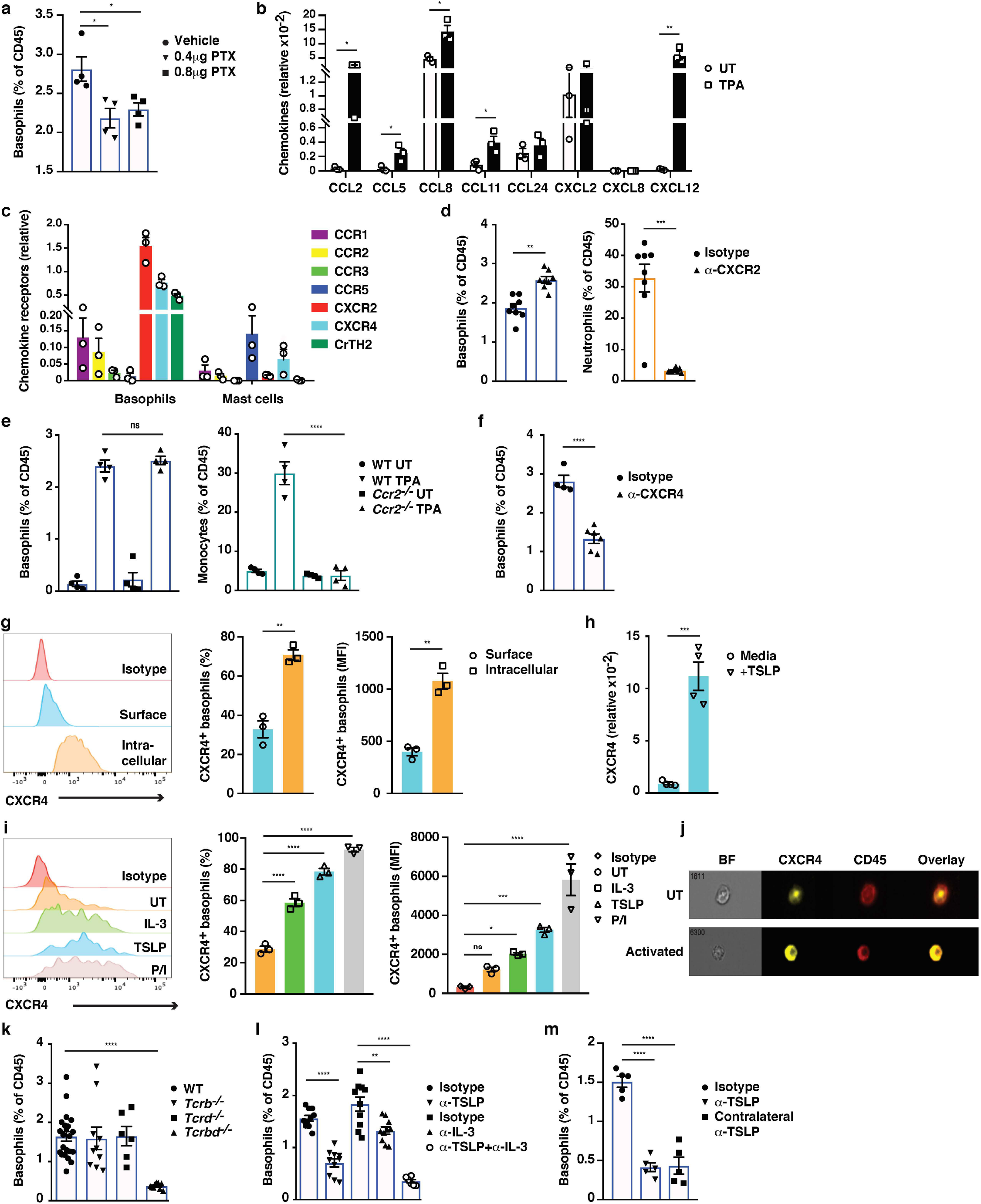
Basophil recruitment to the skin requires TSLP/IL-3-mediated upregulation of CXCR4. (a-f,k-m) Inflamed skin was analysed 24h after a single topical application of 2.5nM TPA and compared to healthy UT skin. (a) Recruitment of basophils to inflamed skin was analysed by FACS and presented as a proportion of all skin CD45_+_ leukocytes following i.v. injection of PTX or vehicle control. Basophils were gated as CD45_lo_cKit_-_IgE_+_CD41_+_. (b) Expression of chemokines in whole skin and (c) chemokine receptors in sorted skin basophils and mast cells was analysed by qRT-PCR and presented relative to the control gene cyclophilin. (d-f) Recruitment of basophils to inflamed skin was analysed as in (a) following: (d) i.v injection of 50ng anti-CXCR2 antibody, (e) in *Ccr2*_*−/−*_ mice or (f) i.v injection of 50ng anti-CXCR4 antibody. Suitable control populations presented to show efficacy of treatment where appropriate. (g-j) Basophils were isolated from the spleen of WT mice and (g) analysed for cell surface or intracellular expression of CXCR4 by flow cytometry, (h) incubated with 1μg/ml TSLP or media alone for 24h and expression of CXCR4 transcripts assessed by qRT-PCR. (i,j) Basophils were activated for 24h with 100ng/ml IL-3, 1μg/ml TSLP or isotype control, with PMA/ionomycin for 4h, or left UT before surface expression of CXCR4 was analysed by (i) FACS or (j) total CXCR4 expression (yellow) imaged with surface staining of CD45 (red) using Imagestream. (k-m) Recruitment of basophils to inflamed skin was analysed as in (a) in (k) *Tcrb*_*−/−*_, *Tcrd*_*−/−*_ and *Tcrbd*_*−/−*_ mice, (l) following i.d. injection of 20ng anti-TSLP or anti-IL-3 (or both) antibodies in the TPA treated ear or (m) in the contralateral ear. Representative data shown in (g,i left panels) as histograms of CXCR4 FACS stain and (j) images from Imagestream, all other data are expressed as means ± SEM. Statistics by one-way ANOVA multiple comparison (a-c,e,g-i and l) or two-tailed Student’s t-test (d,f,j and k); *p<0.05, **p<0.01, ***p<0.001 and ****p<0.0001. ns = not significant. BF = bright field.

### FcεRI-signalling in basophils promotes inflammation-driven outgrowth of cSCCs

As natural IgE promoted EC growth via FcεRI-signalling in basophils during skin inflammation, we next tested whether a similar mechanism may support skin tumour growth. Using the two-stage inflammation-driven model of epithelial carcinogenesis, we found that mice lacking FcεRI (*FceR1a_−/−_*) were significantly less susceptible to tumour development (Fig.7a). Only few mice developed any tumours at all and the tumours that grew were very small (Fig. 7a). This was despite the composition of the immune infiltrate in both the perilesional skin and the tumour tissue was overall similar in *FceR1a_−/−_* and WT mice, apart from a minor reduction in neutrophils in the tumours from *FceR1a*_*−/−*_ mice (Fig. 7b). Further, mice with substantially diminished FcεRI effector cells, *Cpa3*_*Cre/+*_ mice, were also significantly protected from tumour development (Fig. 7c). *Cpa3*_*Cre/+*_ mice lack all mature skin mast cells due to Cre toxicity_17_ and no mast cells were found in neither perilesional skin or tumour tissue (Fig. 7d). However, these mice also have substantially reduced basophil numbers with >90% reduction in inflamed skin and perilesional skin, although basophils were detected in the tumour (Fig. 7d). Due to the dual effect on mast cells and basophils of the *Cpa3*_*Cre*_, we next used *Mcpt8*_*Cre/+*_ mice, which have normal mast cell numbers but a strongly reduced number of basophils (Fig. 7f). The *Mcpt8*_*Cre/+*_ mice were again significantly less susceptible to tumour growth (Fig. 7e), suggesting that FcεRI-signalling in basophils was responsible for tumour growth. As signalling via histamine receptors H_1_R and/or H_4_R on EC were shown to promote EC proliferation and differentiation during skin inflammation, we then tested whether blocking H_1_R affected tumour outgrowth. Indeed, adding Cetirizine, a H_1_R blocker, in the drinking water during tumour promotion significantly delayed tumour onset and suppressed tumour growth (Fig. 7h). Together these data demonstrate that the inflammation-driven tumour promotion is mediated via FcεRI-signalling in skin infiltrating basophils, partly via histamine release.

**Figure 7.**
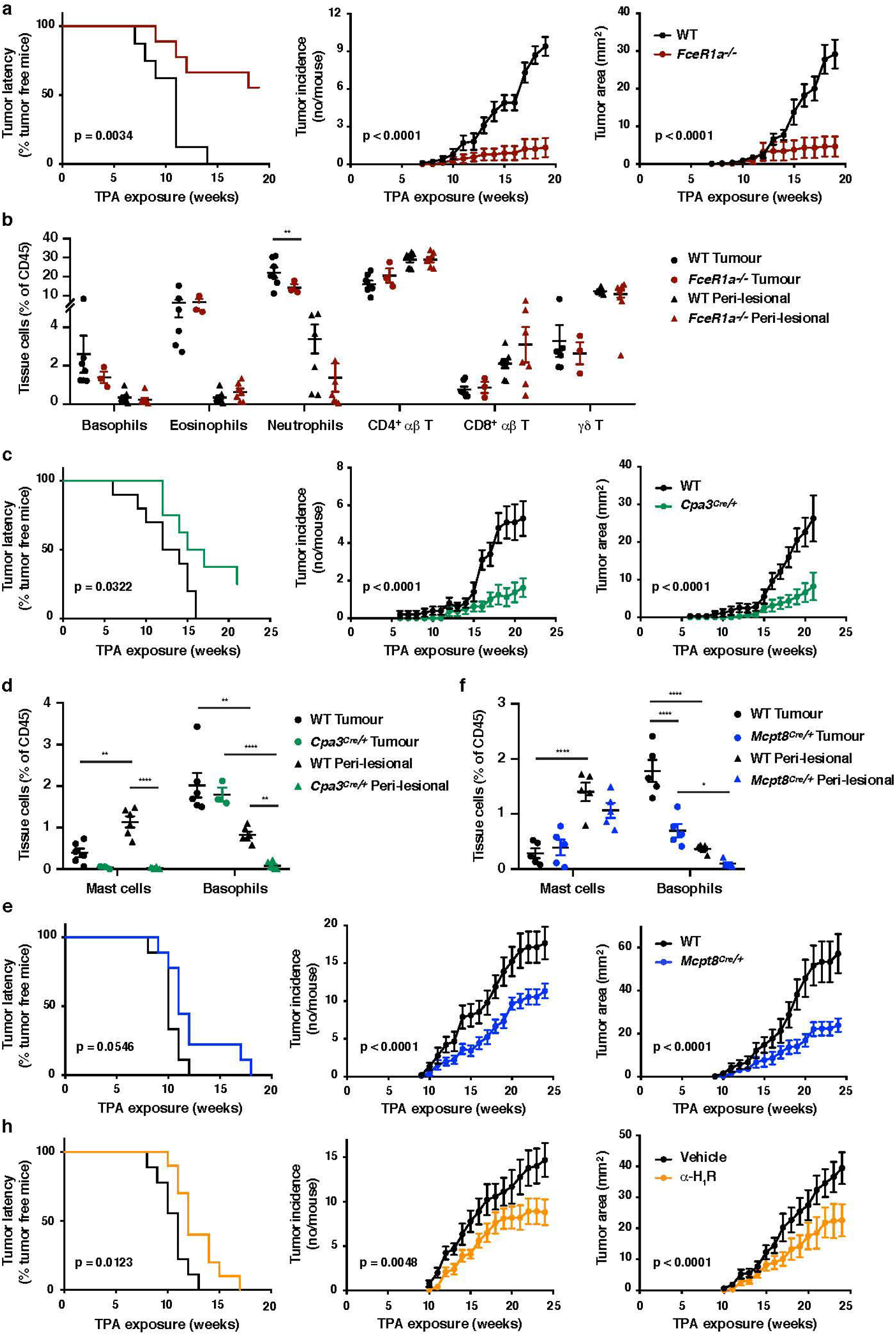
FcεRI-signalling in basophils promote inflammation-driven outgrowth of cSCCs. (a-h) Mice were treated with a single subclinical dose of DMBA to the shaved dorsal back skin and skin inflammation promoted by twice weekly TPA application for 18-25 weeks. Tumor susceptibility is expressed as tumor latency (time to appearance of first tumor), tumor incidence (average number of tumors per mouse) and tumor area (average tumor size per mouse) in (a) WT (n=10) and *FceR1a_−/−_* mice (n=9), (c) WT (n=10) and *Cpa3_Cre/+_* mice (n=8), (e) WT and *Mcpt8*_*Cre/+*_ mice (n=9/group) and (h) WT mice giving H_1_R antagonist Cetirizine at 30 mg/kg or vehicle control in the drinking water throughout the experiment (n=10/group). (b) FACS analysis of leukocyte infiltrate in tumour tissue and perilesional skin of WT and *FceR1a*_*−/−*_ mice (n=7 for WT groups; n=3 for tumour, n=7 for peri-lesional tissue for *FceR1a*_*−/−*_) presented as proportion of total CD45_+_ leukocytes. (d,f) FACS analysis of mast cells and basophils in tumours and peri-lesional tissue of (d) WT and *Cpa3*_*Cre/+*_ mice (n=6 for WT groups; n=3 for tumour, n=6 for peri-lesional tissue for *Cpa3*_*Cre/*_+) and (f) WT and *Mcpt8*_*Cre/+*_ mice (n=5/group). Data are expressed as means ± SEM and statistical significance assessed using Log-rank (Mantel-Cox) test for tumor latency and linear regression for tumor incidence and area (a,c,e,h) and one-way ANOVA multiple comparison (b,d,f); p<0.05, **p<0.01 and ****p<0.0001.

## Discussion

In this study we found that skin inflammation enhanced levels of natural IgE, which accumulated in the tissue mainly on infiltrating basophils recruited via TSLP/IL-3-dependent expression of CXCR4. IgE-activated basophils promoted EC growth and differentiation and strongly drove the outgrowth of inflammation-driven skin SCCs, partly via histamine engagement of H_1_R and H_4_R on EC.

Natural IgE is present at low concentrations in the blood of mice and men but is constitutively available on FcεRI-expressing cells at our barrier surfaces, such as the skin. While the role of natural IgE at steady state has not been widely studied, it is suggested to be part of our ‘frontline’ immune defence against a variety of environmental challenges such as parasitic infections as well as exposure to noxious xenobiotics, irritants and venoms_4, 8_. The host defences evoked may include barrier enhancement, removal or inactivation of the challenge and tissue repair_4, 8_. Our IgE sequencing analysis revealed that IgE induced by low-grade tissue inflammation had a similar repertoire and VDJ gene-usage to that of the natural IgE in healthy animals, suggesting that tissue inflammation merely enhances the level of IgE already present in the host. This IgE fortified the skin barrier defences by inducing EC growth and differentiation and thickening the epidermis. However, this response should only be transiently activated upon exposure to noxious stimuli as long term exacerbation of this response may lead to pathologies and altered EC function. Indeed, we found that when EC harbour oncogenic mutations, chronic activation of the natural IgE response subverts its protective effects and supports the growth of otherwise subclinical EC lesions promoting tumour development.

The tumour-promoting effect of IgE in the context of chronic skin inflammation is in stark contrast to our recently published data showing that IgE provides protection against mutagen-driven EC carcinogenesis_9_. Interestingly, the DNA-damage caused by repeated carcinogen exposure drive a *de novo* induction of autoreactive IgE antibodies, which differ substantially in their repertoire from the inflammation-induced IgE_9_. Repeated carcinogen exposure also gives rise to substantially more mutated tumours than inflammation-driven skin carcinogenesis_18_, suggesting a different mode of tumour outgrowth in the different experimental scenarios. Whole exome sequencing of human cSCC have identified this to be one of the most mutated human cancers with a mean of 50 mutations per megabase of DNA_19_ and also revealed that there are significantly more mutations in immune-related pathways in tumours of high-versus low-risk of metastasis, including in FcεRI-signalling_19_. We also found a significant positive correlation between *FCER1A*-expression in human cSCC and good disease prognosis_9_, which may reflect a broader paradigm as *FCER1A*-expression correlate with overall prolonged survival in many human malignancies_20_. Thus, although the commonly used inflammation-driven cutaneous carcinogenesis mouse model (DMBA-TPA; also used in this study) exhibits recurrent mutations in similar genes (*Hras*, *Kras, Rras2* or *Trp53*) as human cSCC_18_, the low level of mutations may alter the immunogenicity and the microenvironmental niche in which the tumour grows, which may in part explain the paradoxical role of IgE in this model. Together, our previous_9_ and current studies show a potent link between IgE and cancer. However, the biological consequences of IgE engagement clearly depends on the nature of the antibodies and the tissue microenvironment.

Chronic inflammation has frequently been linked to cancer promotion and tumour extrinsic inflammatory conditions may be key to development of selected cancers. Epidemiological studies suggests that low-grade tissue inflammation predisposes to different forms of cancer by contributing to proliferation and survival of malignant cells, promoting angiogenesis and tumour metastasis and by subverting anti-cancer immunity_10, 11, 12_. For example gastric and bladder cancers have since long been known to be driven by inflammation_21, 22, 23_. Interestingly, contrary to many other cancers, RNAseq data in these two cancers suggests a negative correlation between *FCER1A* and overall survival (KMplotter_24_). A recent study also revealed increased mast cell density in human gastric tumours and in the corresponding mouse models, and that mast cell degranulation was driving tumour growth_25_. It should be noted that in this study mast cell numeration was done using toluidine blue in this study, which also stains basophils. Although IgE/FcεRI-signalling has not been linked directly to tumour growth in these cancers, taken together these data suggests that IgE may play a role in cancer promotion in inflammatory settings in some human tissues.

IgE-mediated immune surveillance in the skin is most likely, as in allergic disease, dominated by mast cells in healthy skin and during the early-phase response, whereas the late-phase response is driven by recruited basophils. In our study, basophils were dominant over mast cells in the inflamed skin and only basophils appeared to enter the tumours, whilst mast cells remained at the base of the tumours and in the surrounding tissue. Epidermal hyperplasia during tissue inflammation was driven by FcεRI-signalling in basophils and tumour growth also appeared to be mainly promoted by basophils, although we cannot exclude a role for mast cells as *Cpa3*_*Cre/+*_ mice (lacking mast cells and most basophils) were more protected from carcinogenesis than *Mcpt8*_*Cre/+*_ mice (normal mast cell numbers but lacking most basophils). While the physiological role of basophils remain poorly understood, it is increasingly recognised that basophils regulate immunity at many levels, often distinct from mast cells_7, 26, 27_. For example, skin-infiltrating basophils, but not mast cells, have been shown to be crucial for acquired tick resistance in mice_28_. Interestingly, the mechanism of basophil-mediated tick host defence is via epidermal thickening driven by local histamine release_28_, consistent with the role of basophils and histamine in EC growth demonstrated in our study.

Histamine has wide-ranging effects on skin ECs and has previously been shown to promote wound healing in the skin_29_. For a long time, the effects of histamine in skin diseases have been thought to be mediated solely by its action on H_1_Rs, however newer findings, including this report, suggests H_4_R-signalling to strongly affect skin ECs. H_4_R-deficient mice have thinner epidermis than WT at resting_30_ and H_4_R-deficiency, or blockade, significantly reduces epidermal hyperplasia_31_ and total skin thickness_32_ in inflammatory settings. Surprisingly, mice lacking HDC (the enzyme responsible for histamine generation) are more susceptible to inflammation-induced carcinogenesis of the skin and colon_33_. However, this is due to an important role for histamine in myeloid cell differentiation and maturation within the bone marrow (through H_1_R and H_2_R) and HDC-deficient mice have significantly altered myeloid cell composition including a large increase in immature myeloid cells, which are recruited to the tissue during inflammation and promote cancer growth_33_.

In summary, we demonstrate how IgE antibodies with natural specificities regulate skin EC growth and differentiation via FcεRI-signalling. Further, during chronic skin inflammation IgE strongly promotes outgrowth of ECs harbouring oncogenic mutations. Thus, our data suggest a potent link between autoreactive IgE and skin hyperplasia, which may be pertinent for a variety of inflammatory skin conditions where blocking IgE-mediated signalling could be beneficial, as has been shown for idiopathic urticaria_34, 35_. Our data moreover may have potential important implications for both atopy and cancer.

## Acknowledgements

We thank H.R. Rodewald for providing *Cpa3*_*Cre/+*_ mice. We are indebted to the staff of the Imperial Central Biomedical Services for the care of the animals and the LMS/NIHR Imperial Biomedical Research Centre Flow Cytometry Facility for FACS support. We are grateful to M. Botto and A. Mowat for critical reading of the manuscript and for the informed advice of many close colleagues. This work was supported by the Wellcome Trust (100999/Z/13/Z).

## Author contributions

M.D.H performed and analyzed the experiments with help from S.W., G.C., R.C.S. and W.D.J. G.C. did the antibody sequencing analysis with assistance from D.K. and D.D-W. D.V provided *Mcpt8*_*Cre/+*_ mice and helpful comments. J.S. performed and analyzed some experiments, directed the study and wrote the manuscript.

## Methods

### Mice

Genetically altered mice were generated as previously described. *Igh-7_-/-36_*, *FceR1a_-/-37_*, *Cpa3*_*Cre/+38*_ and *Mcpt8*_*Cre/+39*_ were on the BALB/c background after >10 backcrosses. *Tcrd*_*-/-40*_ and *Tcrb*_*-/-41*_ were on the FVB/N background after > 10 backcrosses and *Tcrdb*_*−/−*_ were generated by intercrossing the respective mutants. *CCR2*_*-/-42*_ mice were on the CD1 background. Non-transgenic littermates were used as controls or strain-matched wild-type control animals were purchased from Charles River. Mice were bred and maintained in individually ventilated cages under specific pathogen-free conditions. Age-matched, female mice were used for all experiments at ≥ 7 weeks of age and selected at random from a large pool when allocated to experiments. All studies were approved by Imperial College AWERB (Animal Welfare and Ethical Review Body) and by the UK Home Office. Experiments involving cancer studies strictly adhered to the guidelines set out by the National Cancer Research Institute (NCRI) and Workman et al. in ‘Guidelines for the Welfare and Use of Animals in Cancer Research’43. All studies using animals were conducted following the Animal Research: Reporting In Vivo Experiments (ARRIVE) guidelines_44_.

### Cutaneous challenge and chemical carcinogenesis

Chemicals 12-0-tetradecanoylphorbol-13-acetate (TPA), MC903 and 7,12-Dimethylbenz[a]anthracene (DMBA) were purchased from Sigma, R848 from Enzo Life Sciences and they were dissolved in ethanol (TPA, MC903) or acetone (DMBA, R848). Skin inflammation was induced by exposing the dorsal sides of the ear skin to a single or repeated dose of either 2.5nM TPA, 1nM MC903 or 100μg R848 in 25μl.

For cutaneous carcinogenesis age-matched female mice were used at 7 weeks of age. The fur of the dorsal back area was removed using hair clippers and mice rested for 1 week. Applications of chemicals and tumor monitoring were performed as previously described_45_. In brief, 600nM DMBA was carefully and slowly applied by pipette, in a 150μl volume, to the entire clipped skin area. Mice were rested for one week and 20nM TPA then applied twice weekly. Hair regrowth during the experiment was gently removed by clipping with trimmers. Mice were monitored daily and cutaneous tumors were counted and measured with a caliper once weekly. Back skin and tumors were evaluated by visual inspection by an observer blinded to the experimental groups.

### Tissue Processing

Skin and tumour tissue were cut into small 1mm_2_ pieces using a scalpel blade and incubated for 2 hrs in digestion buffer containing 25ug/ml Liberase (Roche), 250ug/ml DNAseI (Roche) and 1x DNAse buffer (1.21 Tris base, 0.5g MgCl_2_ and 0.073g CaCl_2_) at 37°. Following digestion, tissue was transferred into C-tubes (Miltenyi Biotech) containing RPMI-1640 media (Thermo Fisher) supplemented with 10% heat-inactivated foetal calf serum, 1% Penicillin-Streptomycin-Glutamine (Thermo Fisher) and physically disrupted using a Miltenyi cell dissociator. Spleen and LN tissue was disaggregated mechanically using a rubber policeman. All cell suspensions were filtered and cells counted using a CASY cell counter (Roche).

### Dermal sheets

Ears were collected, split into dorsal and ventral sides and floated dermis side down in 0.5 M NH4SCN for 40 min, 37 °C and 5% v/v CO2. Subsequently, intact dermal sheets were isolated and washed in PBS. They were then fixed in ice-cold acetone at –20 °C for 15 min, followed by rehydration in PBS and blocking of nonspecific binding with 2% BSA for 1 h. IgE in the tissue was visualized by first incubating with rat anti-mouse IgE (R35-92, BD) followed by Alexa Fluor 555–conjugated goat anti-rat IgE (A21434, ThermoFisher), each for 1 h at room temperature. After extensive washing in PBS, samples were mounted in VectaShield containing DAPI (Vectashield) and visualized with a Leica SP5 confocal laser-scanning microscope (Leica).

### Epidermal thickness

Dorsal side of the ears were treated topically twice a week for 3 weeks with 2.5 nM TPA and subsequently fixed and paraffin embedded. Ear skin from UT mice or the UT contralateral ear was embedded in parallel. 5 um sections were cute and stained for H&E. Images were obtained using a Leica DM4B microscope (Leica). Epidermal thickness was measured using Fiji software by taking 5 measurements per section from 5 sections per ear in a blinded manner. In some experiments H_1_R or H_4_R antagonist was administered by IP injection of 20 mg/kg Cetirizine dihydrochloride (Tocris) or 20 mg/kg JNJ 7777120 (Tocris) respectively or the appropriate vehicle controls.

### Immunofluorescent staining of skin and tumor samples

For ear skin, ears were removed and a defined central section cut. Tumors were removed from the back along with a small piece of adjacent skin. All tissue were snap-frozen in OCT on dry ice. 6-μm sections were cut using a Leica JUNG CM1800 cryostat and stored at –80 °C. For staining, slides were returned to room temperature before fixation with 4% paraformaldhyde for 15 min. Samples were then blocked in 5% goat serum for 1 h at room temperature before staining with rat anti-mouse IgE (R35-92, PharMingen) overnight at 4 °C. Following staining, samples were washed and incubated with Alexa Fluor 555–conjugated goat anti-rat IgG (A21434, ThermoFisher). After extensive washing, samples were then either further incubated with Alexa Flour 647-conjugated rat anti-mouse CD49F (GoH3, BioLegend) or directly mounted in VectaShield containing DAPI (Vectashield). Tissue samples were visualized with a Leica SP5 confocal laser-scanning microscope (Leica).

### Flow cytometry, cell sorting and ImageStream

Cell suspensions were blocked for non-specific binding using antibody against FcγR (2.4G2) and 2% normal rat serum (Sigma) prior to any staining protocols. For staining of cell surface markers, cell suspensions were stained with fluorochrome-conjugated antibodies or appropriate isotype control with the addition of a fixable, live/dead discrimination dye (Invitrogen) for 25 min and subsequently washed. Intracellular staining was then carried out using an Intracellular Staining kit as per manufacturer’s instruction (ThermoFisher Scientific). Cells were fixed for 10 min at 4°C, washed with Perm buffer and stained for intracellular markers for 25 min. For intranuclear γH2AX staining, cells were fixed/permeablized in ice-cold 70% ethanol at −20°C for 2 hrs, blocked with 2% normal mouse serum (Sigma), Fc-block and 2% fetal calf serum for 15 min, followed by 45 min staining for γH2AX at room temperature. Stained cells were analyzed using BD FACSVerse and Fortessa X20 (BD Biosciences, NJ, USA) machines. Data analysis was performed using FlowJo 10 for Mac (TreeStar, OR, USA). For cell sorting a FACS AriaIII High Speed Cell Sorter (BD) was used. For imagestream analysis, internalization scores were determined using CD45 staining as a membrane marker and CXCR4 staining as the probe. Analyses was performed using an ImageStream X Mark II imaging flow cytometer and IDEAS v6 software (AMNIS).

Antibodies were sourced from Biolegend unless otherwise stated. The following antibodies were used: anti-CD11b (M1/70), anti-Siglec-F (E50-2440), anti-Ly6C (HK1.4), anti-Ly6G (IA8), anti-CD4 (GK1.5), anti-CD8 (53-6.7), anti-CXCR4 (I.276F12), anti-CD138 (281-2, BD) anti-CD45 (30-F11, eBioscience), anti-IgE (23G3, eBioscience), anti-TCRβ (H57-597), anti-TCRγδ (eBioGL3, eBioscience), anti-Vγ5 (536, eBioscience), anti-Vγ4 (UC3-10Ab), anti-CD117 (2B8), anti-CD41 (MWReg30), anti-FcεRI (MAR-1), anti-CD95 (Jo2, eBioscience) and γH2AX (JBW301, Millipore). Cells subsets were gated as: Mast cells: CD45_+_CD117_+_IgE/FcεRI_+_CD41_-_; basophils: CD45_mid_CD117_-_IgE/FceRI_+_CD41_+_; eosinophils: CD45_+_GR1_+_CD11b_+_Siglec-F_+_; neutrophils: CD45_+_Ly6G_+_Ly6C_-_CD11b_+_; monocytes/macrophages: CD45_+_CD11b_+_Ly6G_-_Ly6C_+_; αβT cells: CD45_+_Tcrb_+_CD4/CD8_+_; γδ IEL: CD45_+_Tcrd_hi_Vg5_+_; dermal γδ T cell: CD45_+_Tcrd_+_Vg5_-_; plasma cell: FSC_hi_CD95_+_CD138_+_.

### ELISA

Total IgE was measured by an IgE capture method. Sera to be tested and IgE standard were added to plate wells coated with 1 μg/ml rat monoclonal anti-mouse IgE (PharMingen) and blocked with 1% rat serum. Biotinylated rat monoclonal anti-mouse IgE (PharMingen) at 1 μg/ml were then added and incubated for 2h at 37°C. After washing in PBS 0.05% Tween, plates were incubated with alkaline phosphatase-conjugated Streptavidin (PharMingen) for 1 hr at 37°C. After washing, detection of antibody levels was carried out by addition of alkaline phosphatase substrate pNPP (Sigma) and absorbance measured at 405 nm. For detection of anti-phosphorylcholine (anti-PC) specific IgE; total IgE was captured with anti-mouse IgE (PharMingen), followed by addition of PC-BSA at 50 mg/ml (Sigma). After washing, detection was achieved by incubating first with anti-PC IgM antibody (clone BH8) (Millipore), followed by anti-IgM conjugated to alkaline phosphatase (Southern Biotech) and addition of pNPP substrate (Sigma). Absorbance was measured at 405 nm.

For IgG1 and IgG2a antibodies, NUNC Immune Maxisorp 96-well plates (Thermo Scientific) were coated with 5 μg/ml goat anti-mouse IgH+L (Southern Biotech) in borate buffered saline at 37°C for 3hr. After washing, plates were blocked with PBS containing 0.5% BSA for 1 hr at room temperature, and appropriately diluted sera added and incubated overnight at 4°C. After washing, plates were incubated with alkaline phosphatase-conjugated polyclonal goat anti-mouse IgG1 or IgG2a (also detects IgG2c in C57BL/6 mice) (both Southern Biotech) for 5 hr at 4°C. Following further washing, the alkaline phosphatase substrate pNPP (Sigma) was added and absorbance measured at 405 nm.

Histamine levels in serum and ear supernatant were measured using an ultra-sensitive histamine ELISA kit (Enzo Life Sciences) following manufacturer’s instructions. Skin supernatant was acquired by floating the dorsal ear skin on 250 ul of complete media for 16h.

### qRT-PCR and primer sequences

RNA was extracted from isolated cell suspensions or from skin/tumor tissue, preserved in RNA-later, using RNEasy Mini kits (Qiagen). RNA was dissolved in nuclease-free water, and yield and purity were determined. Complementary DNA (cDNA) was synthesised from RNA with an iScript cDNA synthesis kit (Bio-Rad) as per the manufacturer’s instructions. cDNA was diluted in nuclease-free double-deionized water for qRT–PCR. All primers were single-stranded DNA oligonucleotides (Sigma) that were intron-spanning as verified by NCBI Primer-Blast tool. Real-time PCR products were detected with SYBR Green (Life) measured continuously with a ViiA 7 Real-Time PCR system (Applied Biosystems, CA, USA). Ct values for genes of interest were normalized against Ct values of the housekeeping gene Cyclophilin (Cyc) using the 2_-_ΔCt method. The following primers were used: IL-1α: F (5’-TTGGTTAAATGACCTGCAACA-3’), R (5’-GAGCGCTCACGAACAGTTG-3’), IL-4: F (5’-CATCGGCATTTTGAACGAG-3’), R (5’-CGAGCTCACTCTCTGTGGTG-3’), IL-5: F (5’-GAAAGAGACCTTGACACAGCTG-3’), R (5’-GAACTCTTGCAGGTAATCCAGG-3’), IL-6: F (5’-TGATGGATGCTACCAAACTGG-3’), R (5’-TTCATGTACTCCAGGTAGCTATGG-3’), IL-13: (5’-ACACAAGACCAGACTCCCC-3’), R (5’-CTCCTCATTAGAAGGGGCCG-3’), IL-18: F (5’-ACATCTTCTGCAACCTCCAGCA-3’), R (5’-CATTGTTCCTGGGCCAAGAGG-3’), IL-25: (5’-TGGAGCTCTGCATCTGTGTC-3’), R (5’-GATTCAAGTCCCTGTCCAACTC-3’), IL-31: F (5’-GGCCTTCCTCACTCTCTTAC-3’), R (5’-GTATAGGAACCTGGCTGGC-3’), IL-33: F (5’-CACATTGAGCATCCAAGGAA-3’), R (5’-AACAGATTGGTCATTGTATGTACTCAG-3’), TNFα: F (5’-AGCCCACGTAGCAAACCACCA-3’), R (5’-ACACCCATTCCCTTCACAGAGCAAT-3’), Ki67: F (5’-TCTGATGTTAGGTGTTTGAG-3’), R (5’-CACTTTTCTGGTAACTTCTTG-3’), K1: F (5’-TTTGCCTCCTTCATCGACA-3’), R (5’-GTTTTGGGTCCGGGTTGT-3’), K5: F (5’-CCTGCAGAAGGCCAAGCA-3’), R (5’-TGGTGTTCATGAGCTCCTGGTA-3’), K10: F (5’-GGATGAGCTGACCCTTAGCA-3’), R (5’-CATTTTGAAGGTCTCTCATTTCCT-3’), K14: F (5’-CAGCCCCTACTTCAAGACCA-3’), R (5’-GGCTCTCAATCTGCATCTCC-3’), Cpa3: F (5’-CTACGGCCCAATAGCATCCA-3’), R (5’-TGCCCAGGTCATAAACCCAG-3’), Mcpt8: F (5’-CAGTCTATCGCTGTGGTGGT-3’), R (5’-GAGCTTTGCGTTCCAGCTTC-3’), HDC: F (5’-GAGTGCACAGCACAGACAAAGG-3’), R (5’-TCTAGCTCGGTAGTATTCACT-3’), Ptgds: F (5’-GACACAGTGCAGCCCAACTTTC-3’), R (5’-GGGCTACCACTGTCTTGCACATA-3’),Hptgds: F (5’-ATCCACCAGAGCCTCGCAATAG-3’), R (5’-TCATCCAGCGTGTCCACCA-3’), Ptges1: F (5’-GGATGCGCTGAAACGTGGA-3’), R (5’-CAGGAATGAGTACACGAAGCC-3’), Ptges2: F (5’-CTCATCAGCAAGCGCCTCAA-3’), R (5’-GGTCTTTACCCACGGCTGTCA-3’), Ptges3: F (5’-ATCACATGGGTGGTGATGAGGA-3’), R (5’-AGGCGATGACAACAGCCCTTAC-3’), CCL2: F (5’-CCCAATGAGTAGGCTGGAGA-3’), R (5’-AAAATGGATCCACACCTTGC-3’), CCL5: F (5’-TGCCCACGTCAAGGAGTATTTC-3’), R (5’-AACCCACTTCTTCTCTGGGTTG-3’), CCL8: F (5’-CCCTTCGGGTGCTGAAAAG-3’), R (5’-TCTGGAAAACCACAGCTTCCA-3’), CCL11: F (5’-ATGCACCCTGAAAGCCATAGTC-3’), R (5’-CAGGTGCTTTGTGGCATCCT-3’), CCL24: F (5’-CGGCCTCCTTCTCCTGGTA-3’), R (5’-TGGCCAACTGGTAGCTAACCA-3’), CXCL2: F (5’-CGCTGTCAATGCCTGAAG-3’), R (5’-GGCGTCACACTCAAGCTCT-3’), CXCL8: F (5’-CTCTTGGCAGCCTTCCTGATT-3’), R (5’-TATGCACTGACATCTAAGTTCTTTAGCA-3’), CXCL12: F (5’-GAGCCAACGTCAAGCATCTG-3’), R (5’-CGGGTCAATGCACACTTGTC-3’), CCR1: F (5’-AAGGCCCAGAAACAAAGTCT-3’), R (5’-TCTGTAGTTGTGGGGTAGGC-3’), CCR2: F (5’-ACACCCTGTTTCGCTGTAGG-3’), R (5’-GATTCCTGGAAGGTGGTCAA-3’), CCR3: F (5’-ATGGCATGTGTAAGCTCCTCTCAG-3’), R (5’-TTGCTCCGCTCACAGTCATTTCCCCR-3’), CXCR2: F (5’-AGCAAACACCTCTACTACCCTCTA-3’), R (5’-GGGCTGCATCAATTCAAATACCA-3’), CXCR4: F (5’-TCAACCTCTACAGCAGCGTTCTCTT-3’), R (5’-TGTTGGTGGCGTGGACAAT-3’), CrTH2: F (5’-TCTCAACCAATCAGCACACC-3’), R (5’-CCTCCAAGAGTGGACAGAGC-3’), Cyc: F (5’-CAAATGCTGGACCAAACACAA-3’), R (5’-CCATCCAGCCATTCAGTCTTG-3’).

### Immunoglobulin sequencing and analysis

Mice were treated with TPA to the dorsal side of the ear skin twice a week for 2 weeks. 48h after the last exposure, draining-LNs were collected and plasma cells (FSC_hi_CD95_hi_CD138_+_) cell sorted on a BD FACS Aria III (BD Biosciences, NJ, USA). RNA was extracted using RNEasy Micro kits (Qiagen) and reverse transcriped using SuperScript™ III Reverse Transcriptase (ThermoFisher). RNA was also collected from whole spleens of naïve mice and reverse transcriped similarly. Sequences were amplified using Phusion High-Fidelity DNA Polymerase (NewEngland Biolabs) to ensure accuracy and robust performance for large PCR products. PCR with a primer in the constant Cε region 5’-CTAGGGTCATGGAAGCAGTGCC-3’ in combination with a promiscuous V region primer 5’-GAGGTGCAGCTGCAGGAGTCTGG-3’ was performed. Primers were labeled at either end with Multiplex Identifiers (MIDs) for multiplexing. PCR thermal cycling was as follows: 98°C, 30 sec; 30x (98°C, 30 sec; 60°C, 30 sec; 72°C, 35 sec); and 72°C, 10min. Amplicons were purified by gel extraction using QIAquick PCR purification kit (Qiagen) and deep-sequenced by long-read 454 pyrosequencing on the Genome Sequencer FLX system (Roche). Raw sequencing data was presented in. FASTA format and unproductive sequences removed (data cleanup stages as described_46_).

Analysis was performed as previously described_46_. In short: sequences were assigned to individual samples according to their MID. V(D)J gene assignment of individual sequences were annotated using IMGT/HighV-Quest and the mouse database. Physiochemical properties of the CDRH3 region were calculated using the R package Peptides. Physiochemical properties included length, isoelectric point (pI) and frequencies of amino acid classes in the CDRH3 region.

### Bone marrow-derived basophils and mast cells

Bone marrow was obtained from tibias and femurs of 8-10 week old BALB/c mice and erythrocytes depleted using Lysis Buffer Hybri-MaxTM (Sigma). Cells were washed and seeded in T25 flasks (TPP Tissue Culture Flasks, Sigma) in 10 ml RPMI supplemented with 10% heat inactivated foetal bovine serum (Gibco, Life Technologies), 1x penicillin streptomycin glutamine (PSG) (Gibco), 0.1 mM 2-Mercaptoethanol (Gibco) and 10 ng/mL rIL-3 (PeproTech, London, UK). Cells were incubated at 37°C, 5% CO2 in a humidified incubator and the media refreshed every 5 days. Basophil numbers would peak after ~14 days and mast cells after ~1 month. For some experiments, cells were loaded with purified mouse IgE antibodies (R35-92, BD) at 0.5 μg/mL for 24 hours and subsequently crosslinked with anti-IgE (R35-72, BD) at 5 μg/mL. IgE-crosslinked cells were harvested for qRT-PCR analyses and the supernatants stored.

### Primary neonatal keratinocyte cultures

Total body wall skin from neonatal mice (<5 days old) was incubated overnight at 4 °C in 5 U/ml Dispase (BD) supplemented with 1x antibiotic and antimycotic solution (Sigma). The epidermis was isolated and further digested in TrypLE Express supplemented with 200 μg/ml DNAse I and DNAse buffer. Cell suspensions were filtered and resuspended in defined KC serum-free medium with supplements (Life) and 1x antibiotic–antimycotic solution. Keratinocytes were seeded at an appropriate cell density onto tissue culture vessels coated with rat tail-derived collagen I (Sigma). Culture vessels were washed with PBS 24 h following seeding to remove unattached cells, and provided with fresh medium ± addition of histamine (Sigma) or conditioned media from IgE-crosslinked basophils. In some experiments histamine receptors were blocked using 10 μM H_1_R antagonist Cetirizine dihydrochloride or H_4_R antagonist JNJ 7777120 (both from Tocris).

### Cellular recruitment to the skin

To block cell recruitment to the skin, intradermal injections of 20 ng anti-TSLP, 20 ng anti-IL-3, both or isotype controls were injected into the dorsal ear pinnae. In other experiments mice received IV injections of either 0.4 ug PTX or 0.8 ug PTX (Tocris), 50 ng anti-CXCR2 (R&D), 50 ng anti-CXCR4 (R&D) or suitable controls. Skin inflammation was initiated using a single dose of 2.5 nM of TPA to the dorsal ear skin and the skin analysed 24h later.

### Statistical evaluation

The statistical significance of difference between experimental groups was determined using two-tailed Student’s t-test for unpaired data, one-way ANOVA multiple comparison, Log-rank Mantel-Cox test or linear regression, where appropriate, with results deemed significant at p<0.05. Stars of significance correlate to: *p<0.05; **P<0.01; ***P<0.001 and ****p<0.0001. Statistics was performed with GraphPad Prism 6.00 for Mac (GraphPad; La Jolla, CA, USA).

### Data availability

The data supporting the findings of this study are available from the corresponding author upon request. RNA sequencing data is available from the public repository on the National Center for Biotechnology Information’s Sequence Read Archive in raw format (BioProject: PRJNA417372; BioSample accession: SAMN07985450, SAMN07985451, SAMN07985452, SAMN07985453, SAMN07985454, SAMN07985455).

---

**Supplementary figure 1. IgE-deficient mice acquire DNA-damage when exposed to DMBA but develop less antibodies during DMBA-TPA carcinogenesis**

(a) WT and *Igh7*_*−/−*_ mice were treated with a single dose of 200nmol DMBA to the dorsal ear skin and phosphorylation of the histone H2AX (γH2AX), as a measure of double-stranded DNA-breaks, were measured by FACS in CD45_-_ skin epithelial cells 24h later (n=6 for DMBA-treated WT, n=5 in all other groups). (b) WT and *Igh7*_*−/−*_ mice were treated with a single subclinical dose of DMBA to the shaved dorsal back skin and skin inflammation promoted by twice weekly TPA application for 19 weeks. Total IgE, IgG1and IgG2a was measured in the serum by ELISA at end point (n=9/group). Data are expressed as means ± SEM and statistical significance assessed by one-way ANOVA multiple comparison (a) or two-tailed Student’s t-test for unpaired data (b); **p<0.01 and ***p<0.001. ns = not significant, nd = not detected.

**Supplementary figure 2. The cellular immune composition of inflamed skin is unaltered in the absence of IgE-mediated signalling**

(a,b) Ear skin of WT and (a) *Igh7_−/−_ or (b) FceR1a*_*−/−*_ mice were treated topically with TPA, twice a week for two weeks, and the cellular immune composition of the skin analysed by FACS and compared to UT healthy skin. Data are presented as means ± SEM of total CD45_+_ leukocytes in the skin. Gating strategy for specific cell populations is outlined in methods. Statistics by one-way ANOVA multiple comparison; *p<0.05, **p<0.01 and ****p<0.0001.

**Supplementary figure 3. IgE-activated basophils promote cytokine expression in skin EC**

Neonatal WT skin EC (keratinocytes) were grown *in vitro* to 70% confluency and then supplemented with media from IgE-crosslinked basophils or media alone (n=3/group). EC expression of indicated cytokines were analysed by qRT-PCR and expressed relative to the control gene cyclophilin. Data are presented as means ± SEM and statistical significance assessed by two-tailed Student’s t-test; *p<0.05 and ***p<0.001.

## References

1. Geha, R.S., Jabara, H.H. & Brodeur, S.R. The regulation of immunoglobulin E class-switch recombination. Nat Rev Immunol 3, 721–732 (2003).

2. MacGlashan, D., Jr. IgE receptor and signal transduction in mast cells and basophils. Curr Opin Immunol 20, 717–723 (2008).

3. McCoy, K.D. et al. Natural IgE production in the absence of MHC Class II cognate help. Immunity 24, 329–339 (2006).

4. Palm, N.W., Rosenstein, R.K. & Medzhitov, R. Allergic host defences. Nature 484, 465–472 (2012).

5. Lawrence, M.G. et al. Half-life of IgE in serum and skin: Consequences for anti-IgE therapy in patients with allergic disease. J Allergy Clin Immunol 139, 422–428 e424 (2017).

6. Crivellato, E., Beltrami, C., Mallardi, F. & Ribatti, D. Paul Ehrlich’s doctoral thesis: a milestone in the study of mast cells. Br J Haematol 123, 19–21 (2003).

7. Voehringer, D. Protective and pathological roles of mast cells and basophils. Nat Rev Immunol 13, 362–375 (2013).

8. Profet, M. The function of allergy: immunological defense against toxins. Q Rev Biol 66, 23–62 (1991).

9. Crawford, G. et al. Epithelial damage and tissue gammadelta T cells promote a unique tumor-protective IgE response. Nat Immunol 19, 859–870 (2018).

10. Mantovani, A., Allavena, P., Sica, A. & Balkwill, F. Cancer-related inflammation. Nature 454, 436–444 (2008).

11. Coussens, L.M., Zitvogel, L. & Palucka, A.K. Neutralizing tumor-promoting chronic inflammation: a magic bullet. Science 339, 286–291 (2013).

12. Grivennikov, S.I., Greten, F.R. & Karin, M. Immunity, Inflammation, and Cancer. Cell 140, 883–899 (2010).

13. Kearney, J.F. Innate-like B cells. Springer Seminars in Immunopathology 26, 377–383 (2005).

14. Abel, E.L., Angel, J.M., Kiguchi, K. & DiGiovanni, J. Multi-stage chemical carcinogenesis in mouse skin: fundamentals and applications. Nat Protoc 4, 1350–1362 (2009).

15. Ohnmacht, C. et al. Basophils Orchestrate Chronic Allergic Dermatitis and Protective Immunity against Helminths. Immunity 33, 364–374 (2010).

16. Dalessandri, T. & Strid, J. Beneficial autoimmunity at body surfaces - immune surveillance and rapid type 2 immunity regulate tissue homeostasis and cancer. Front Immunol 5, 347 (2014).

17. Feyerabend, T.B. et al. Cre-mediated cell ablation contests mast cell contribution in models of antibody- and T cell-mediated autoimmunity. Immunity 35, 832–844 (2011).

18. Nassar, D., Latil, M., Boeckx, B., Lambrechts, D. & Blanpain, C. Genomic landscape of carcinogen-induced and genetically induced mouse skin squamous cell carcinoma. Nat Med 21, 946–954 (2015).

19. Inman, G.J. et al. The genomic landscape of cutaneous SCC reveals drivers and a novel azathioprine associated mutational signature. Nature Communications 9 (2018).

20. Gentles, A.J. et al. The prognostic landscape of genes and infiltrating immune cells across human cancers. Nat Med 21, 938–945 (2015).

21. Fox, J.G. & Wang, T.C. Inflammation, atrophy, and gastric cancer. Journal of Clinical Investigation 117, 60–69 (2007).

22. Echizen, K. et al. NF-kappa B-induced NOX1 activation promotes gastric tumorigenesis through the expansion of SOX2-positive epithelial cells. Oncogene 38, 4250–4263 (2019).

23. Michaud, D.S. Chronic inflammation and bladder cancer. Urologic Oncology-Seminars and Original Investigations 25, 260–268 (2007).

24. Nagy, A., Lanczky, A., Menyhart, O. & Gyorffy, B. Validation of miRNA prognostic power in hepatocellular carcinoma using expression data of independent datasets. Scientific Reports 8 (2018).

25. Eissmann, M.F. et al. IL-33-mediated mast cell activation promotes gastric cancer through macrophage mobilization. Nature Communications 10 (2019).

26. Karasuyama, H., Mukai, K., Tsujimura, Y. & Obata, K. Newly discovered roles for basophils: a neglected minority gains new respect. Nature Reviews Immunology 9, 9–13 (2009).

27. Schwartz, C., Eberle, J.U. & Voehringer, D. Basophils in inflammation. European Journal of Pharmacology 778, 90–95 (2016).

28. Tabakawa, Y. et al. Histamine Released From Skin-Infiltrating Basophils but Not Mast Cells Is Crucial for Acquired Tick Resistance in Mice. Frontiers in Immunology 9 (2018).

29. Gutowska-Owsiak, D., Selvakumar, T.A., Salimi, M., Taylor, S. & Ogg, G.S. Histamine enhances keratinocyte-mediated resolution of inflammation by promoting wound healing and response to infection. Clinical and Experimental Dermatology 39, 187–195 (2014).

30. Glatzer, F. et al. Histamine induces proliferation in keratinocytes from patients with atopic dermatitis through the histamine 4 receptor. Journal of Allergy and Clinical Immunology 132, 1358–1367 (2013).

31. Rossbach, K. et al. Histamine H4 receptor knockout mice display reduced inflammation in a chronic model of atopic dermatitis. Allergy 71, 189–197 (2016).

32. Cowden, J.M., Zhang, M., Dunford, P.J. & Thurmond, R.L. The Histamine H-4 Receptor Mediates Inflammation and Pruritus in Th2-Dependent Dermal Inflammation. Journal of Investigative Dermatology 130, 1023–1033 (2010).

33. Yang, X.D. et al. Histamine deficiency promotes inflammation-associated carcinogenesis through reduced myeloid maturation and accumulation of CD11b(+)Ly6G(+) immature myeloid cells. Nature Medicine 17, 87–U263 (2011).

34. Maurer, M. et al. Efficacy and safety of omalizumab in patients with chronic urticaria who exhibit IgE against thyroperoxidase. Journal of Allergy and Clinical Immunology 128, 202–U326 (2011).

35. Maurer, M. et al. Omalizumab for the Treatment of Chronic Idiopathic or Spontaneous Urticaria. New England Journal of Medicine 368, 924–935 (2013).

## Method references

36. Oettgen, H.C. et al. Active anaphylaxis in IgE-deficient mice. Nature 370, 367–370 (1994).

37. Dombrowicz, D., Flamand, V., Brigman, K.K., Koller, B.H. & Kinet, J.P. Abolition of anaphylaxis by targeted disruption of the high affinity immunoglobulin E receptor alpha chain gene. Cell 75, 969–976 (1993).

38. Feyerabend, T.B. et al. Cre-mediated cell ablation contests mast cell contribution in models of antibody- and T cell-mediated autoimmunity. Immunity 35, 832–844 (2011).

39. Ohnmacht, C. et al. Basophils Orchestrate Chronic Allergic Dermatitis and Protective Immunity against Helminths. Immunity 33, 364–374 (2010).

40. Itohara, S. et al. T cell receptor delta gene mutant mice: independent generation of alpha beta T cells and programmed rearrangements of gamma delta TCR genes. Cell 72, 337–348 (1993).

41. Mombaerts, P., Clarke, A.R., Hooper, M.L. & Tonegawa, S. Creation of a large genomic deletion at the T-cell antigen receptor beta-subunit locus in mouse embryonic stem cells by gene targeting. Proc Natl Acad Sci U S A 88, 3084–3087 (1991).

42. Boring, L. et al. Impaired monocyte migration and reduced type 1 (Th1) cytokine responses in C-C chemokine receptor 2 knockout mice. Journal of Clinical Investigation 100, 2552–2561 (1997).

43. Workman, P. et al. Guidelines for the welfare and use of animals in cancer research. Br J Cancer 102, 1555–1577 (2010).

44. Kilkenny, C., Browne, W.J., Cuthill, I.C., Emerson, M. & Altman, D.G. Improving bioscience research reporting: the ARRIVE guidelines for reporting animal research. PLoS Biol 8, e1000412 (2010).

45. Strid, J. et al. Acute upregulation of an NKG2D ligand promotes rapid reorganization of a local immune compartment with pleiotropic effects on carcinogenesis. Nat Immunol 9, 146–154 (2008).

46. Wu, Y.C., Kipling, D. & Dunn-Walters, D. Assessment of B Cell Repertoire in Humans. Methods Mol Biol 1343, 199–218 (2015).

