## Supplementary figures and images for "Natural IgE promotes epithelial hyperplasia and inflammation-driven tumour growth"

### Supplemental Figure 1

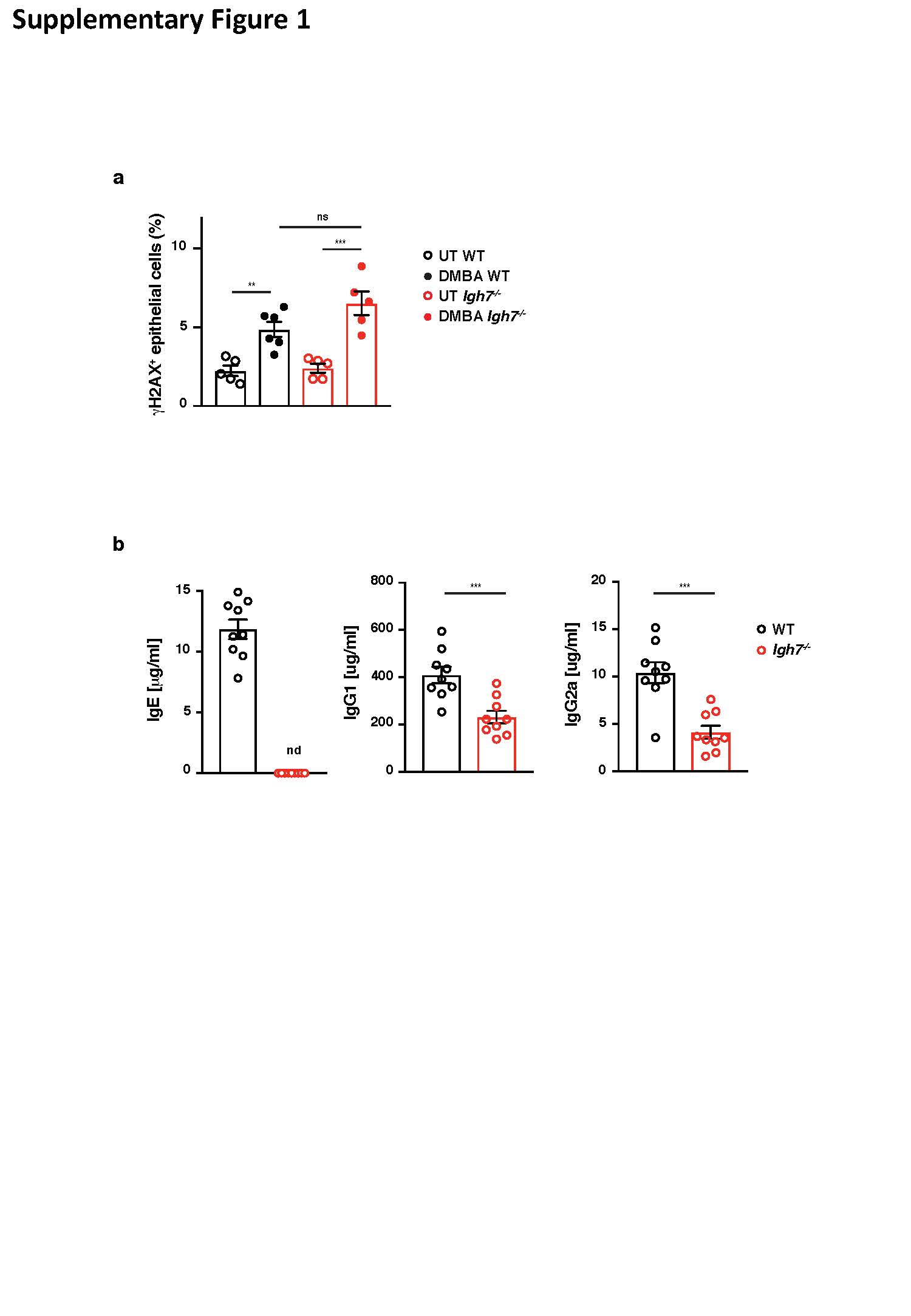

### Supplemental Figure 2

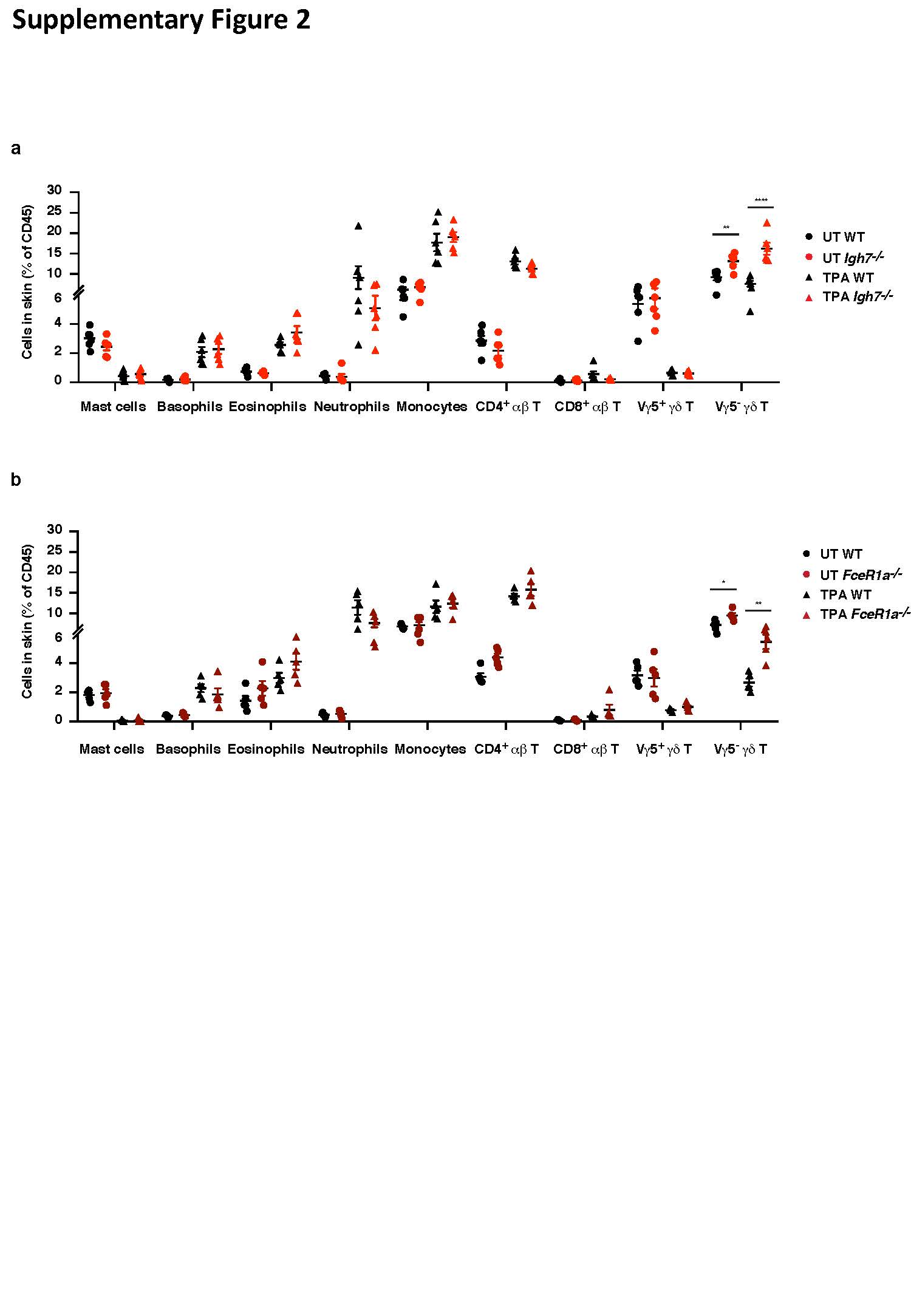

### Supplemental Figure 3

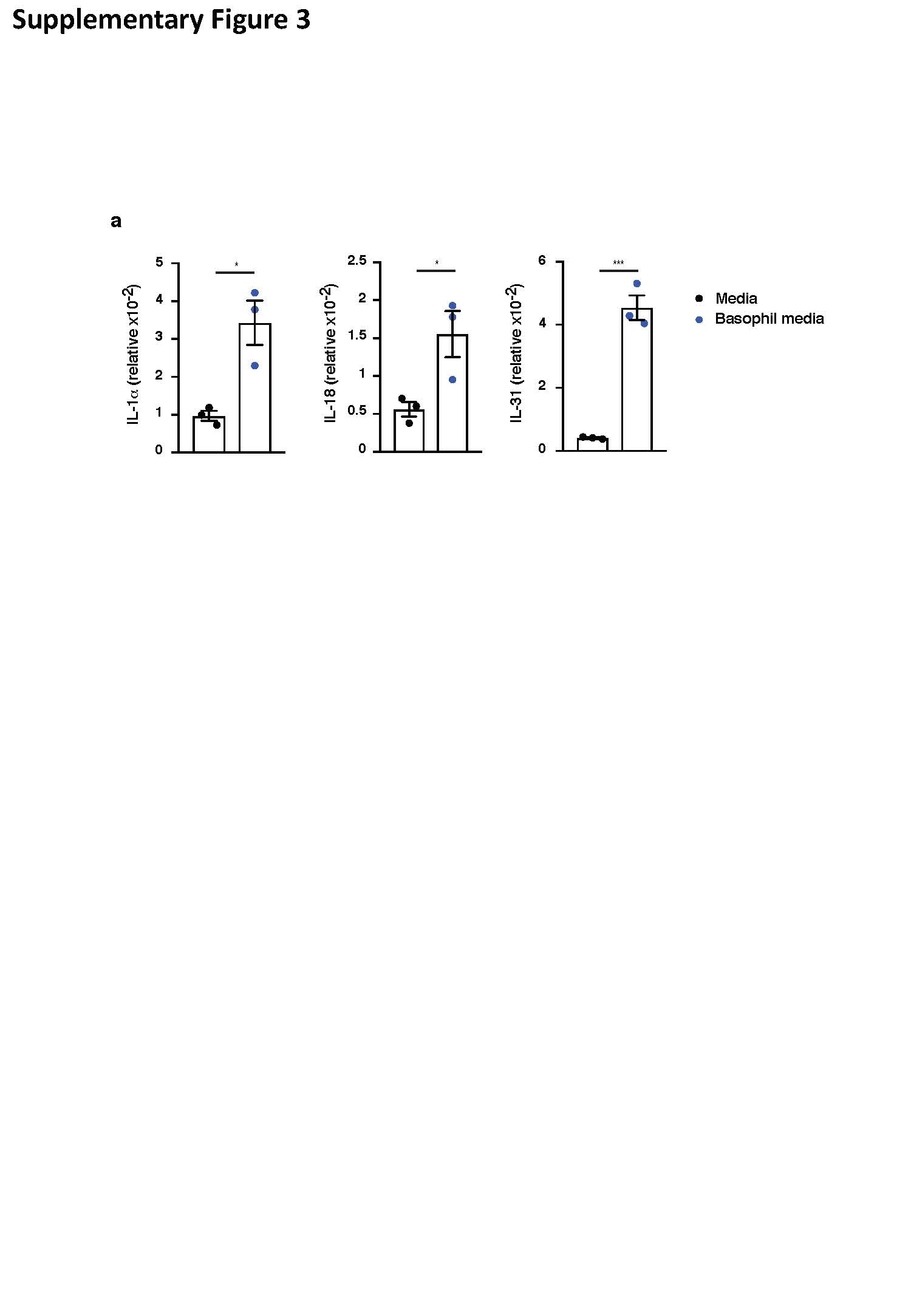
